# Architecture of Ca^2+^ tunneling, a basic Ca^2+^ signaling modality important for secretion

**DOI:** 10.1101/2023.12.17.572039

**Authors:** Raphael J. Courjaret, Larry E. Wagner, Rahaf R. Ammouri, Lama Assaf, Fang Yu, Melanie Fisher, Mark Terasaki, David I. Yule, Khaled Machaca

## Abstract

Ca^2+^ tunneling is a signaling modality that requires both Store-operated Ca^2+^ entry (SOCE) and Ca^2+^ release from the endoplasmic reticulum (ER). Tunneling expands the SOCE microdomain at ER-plasma membrane (PM) contact sites (ERPMCS) through Ca^2+^ uptake by the sarco/endoplasmic reticulum Ca^2+^ ATPase (SERCA) into the ER lumen where it diffuses and is released via open inositol trisphosphate (IP_3_) receptors (IP_3_Rs). In this study using high resolution imaging, we outline the spatial remodeling of the Ca^2+^ tunneling machinery (IP_3_R1; SERCA; PMCA; and Ano1 as an effector) relative to STIM1 in response to store depletion. We show that store depletion leads to redistribution of these Ca^2+^ signaling modulators to distinct subdomains laterally at the PM and axially within the cortical ER. To functionally define the role of Ca^2+^ tunneling, we engineered a Ca^2+^ tunneling attenuator (CaTAr) that blocks tunneling without affecting Ca^2+^ release or SOCE. CaTAr inhibits Cl^−^ secretion in sweat gland cells. Viral mediated expression of CaTAr in the mouse reduces sweating, showing that Ca^2+^ tunneling is important physiologically. Collectively our findings outline the architecture of the Ca^2+^ tunneling machinery and show that it is a fundamental physiological pertinent Ca^2+^ signaling modality.

## INTRODUCTION

Agonists activate cell surface receptors triggering signaling cascades that allow the cell to respond to environmental cues and coordinate with other cells and tissues within the organism to maintain homeostasis. Ca^2+^ signaling is a primary signaling modality downstream of receptors linked to the activation of phospholipase C (PLC), which hydrolyzes the membrane lipid phosphatidylinositol 4,5-bisphosphate (PIP2) producing Inositol 1,4,5-trisphosphate (IP_3_) and diacylglycerol (DAG) ^1^. IP_3_ binds to the IP_3_ receptor (IP_3_R), which is an ER Ca^2+^ permeable channel, resulting in a transient Ca^2+^ release phase as store Ca^2+^ content is limited. Depletion of ER Ca^2+^ stores activates Store operated Ca^2+^ entry (SOCE) leading to a smaller more sustained Ca^2+^ influx phase from the extracellular space ^2^. Ca^2+^ release and SOCE are coupled through a third Ca^2+^ signaling modality known as Ca^2+^ tunneling (Fig. 1A). During tunneling, Ca^2+^ entering the cell through SOCE channels is taken up by the sarco/endoplasmic reticulum (ER) Ca^2+^ ATPase (SERCA) into the ER and released through IP_3_Rs ^3–5^. Tunneling spatially expands SOCE signaling as Ca^2+^ diffuses more efficiently within the ER lumen due its lower Ca^2+^ buffering capacity compared to the cytosol ^6–10^. Tunneling also modulates the spatial, temporal, and oscillation dynamics of the Ca^2+^ signal ^3–5, 11, 12^. Ca^2+^ tunneling was originally described in pancreatic acinar cells ^4, 13, 14^, and has more recently has been demonstrated in oocytes and HeLa cells, where the tunneled Ca^2+^ signal is mostly cortical and does not reach effectors deep within the cell (Fig. 1A) ^11, 12, 15^.

**Figure 1:**
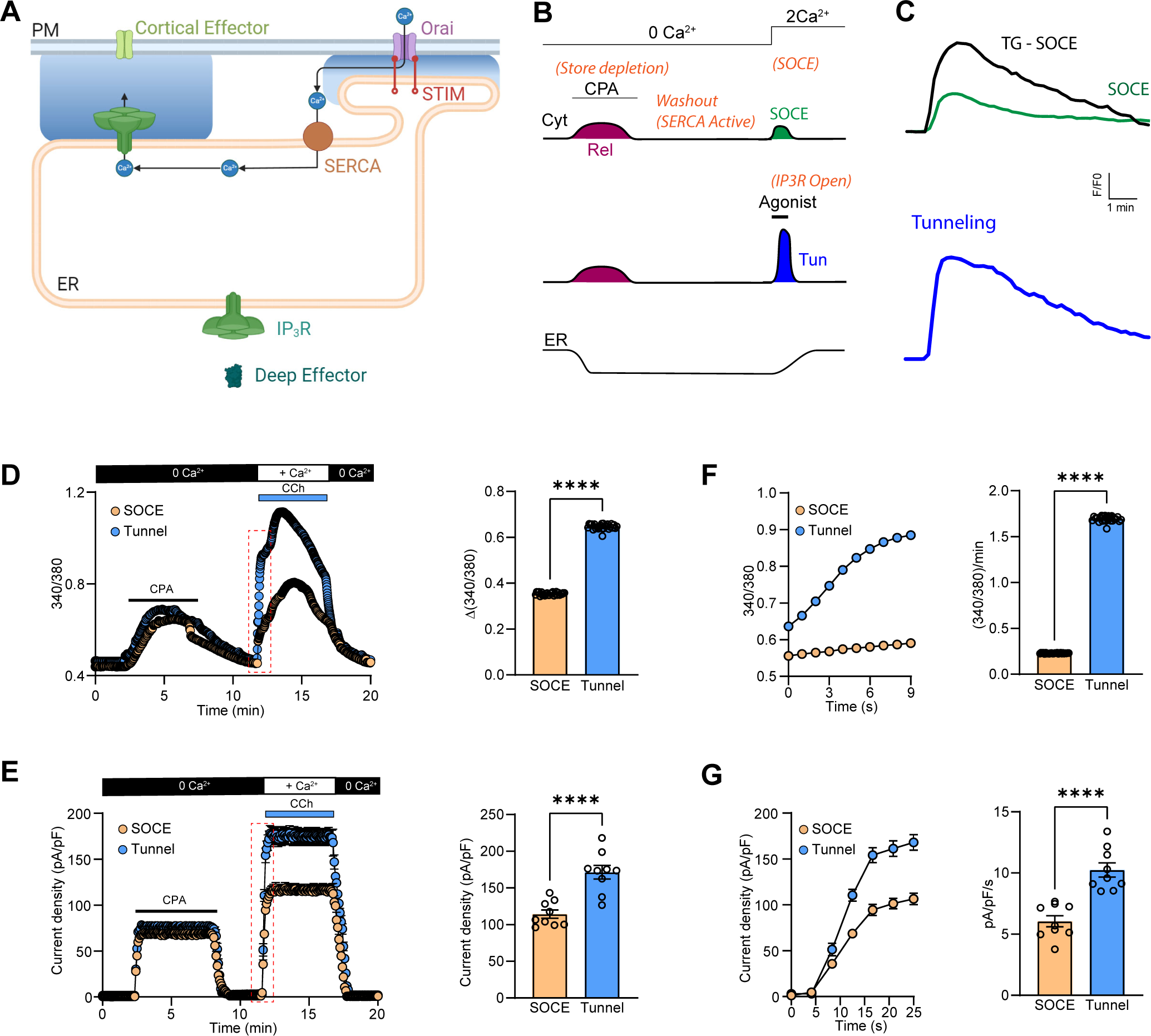
Ca^2+^ Tunneling in primary salivary gland cells. (**A**) Cartoon depiction of Ca^2+^ tunneling. STIM1-Orai1 generate localized Ca^2+^ influx within ERPMCS defining the SOCE microdomain. Ca^2+^ taken into the ER by SERCA diffuses through the cortical ER and is released by IP_3_R to activate distal cortical targets. **(B)** Comparison of SOCE and Ca^2+^ tunneling protocols. The ER Ca^2+^ stores are depleted by the reversible SERCA inhibitor CPA. This is followed by a washout phase to restore SERCA activity. SOCE develops upon addition of extracellular Ca^2+^. Concurrent exposure to an agonist to produce IP_3_ activates Ca^2+^ tunneling. **(C)** Example traces of the intracellular Ca^2+^ signals recorded at the PM during SOCE, SOCE induced by the irreversible blockade of the SERCA pump by thapsigargin (SOCE-Tg) and during tunneling. Traces are taken from Courjaret et al. 2018 ^11^. **(D)** Cytosolic Ca^2+^ responses during SOCE and tunneling in isolated salivary gland cells. Tunneling is induced by the addition of CCh (10 µM) simultaneously to Ca^2+^ reperfusion. The bar chart quantifies the peak Ca^2+^ signal of tunneling *vs* SOCE (n=26; unpaired t-test). **(E)** Whole cell Cl^−^ currents in response to SOCE vs tunneling, showing similar profiles to the Ca^2+^ signals (n=9; unpaired t-test). **(F, G)** Kinetics of intracellular Ca^2+^ elevation and Cl^−^ current development during tunneling compared to SOCE (n=26 (F) and 9 (G); unpaired t-test).

SOCE is mediated by two classes of proteins; the STIM resident ER Ca^2+^ sensors (STIM1, STIM2) with lumenal Ca^2+^ binding domains, and the Orai Ca^2+^-selective PM channels (Orai1, Orai2, Orai3). Store depletion induces a conformational change in STIM1 resulting in its clustering into higher order oligomers and exposing the SOAR/CAD domain, which directly binds to and gates Orai1 ^16–19^. Clustered STIM1 is enriched at ER-PM contact sites (ERPMCS) where it recruits Orai1 by diffusional trapping, and gates it open thus activating SOCE ^20–22^. SOCE is critical for immune cell activation, muscle development, and secretion ^23^. This is highlighted by the defects observed in patients with mutations in either STIM1 or Orai1-which phenocopy each other-, including severe combined immunodeficiency, muscle hypotonia, and ectodermal dysplasia, including defective sweating revealed as anhidrosis with dry skin and heat intolerance ^24, 25^.

Given the requirement for a direct physical interaction between STIM1 and Orai1 for SOCE, their colocalization to ERPMCS is essential. ERPMCS are close appositions between the ER and PM where the two membranes are <30 nm apart allowing the STIM1 cytoplasmic domain in its extended conformation to span the gap and activate Orai1 ^26–28^. The gap distance between the ER and PM is controlled by different tethering proteins physiologically and can be modulated experimentally using artificial linkers ^29–32^. ERPMCS are present at steady state (when Ca^2+^ stores are full) and are stabilized by tethering proteins such as the extended synaptotagmins (E-Syt), TMEM24, ORPs, and GRAMDs ^33^. Store depletion leads to lateral expansion of ERPMCS ^31, 34^. Because ERPMCS are the sites of SOCE their physical dimensions and distribution are critical to understand SOCE signaling and Ca^2+^ tunneling.

The structure of the SOCE microdomain at ERPMCS has been comprehensively discussed by Hogan ^35^. The lateral spread of endogenous ERPMCS estimated from electron microscopy studies ranges from 100-300 nm and they occupy 1-4 % of the total PM area ^26, 27, 35^. This limited footprint is coupled to a relatively small number of Orai1 channels within ERPMCS after store depletion, estimated at 1-5 channels with a predicted single channel current of ∼2 fA and a probability of opening (P_O_) of ∼0.8 ^35, 36^. These estimates argue that SOCE Ca^2+^ signals are spatially confined within ERPMCS to a small percent of the cell cortex. Furthermore, Ca^2+^ diffusion out of the SOCE microdomain at ERPMCS is limited by the high cytosolic Ca^2+^ buffering capacity ^6^, as well as Ca^2+^ uptake and extrusion by Ca^2+^ pumps at the PM (PMCA) and at the ER (SERCA). Measurements of the cortical Ca^2+^ signal due to SOCE alone (when SERCA is active) shows a transient low amplitude signal (Fig. 1B and 1C, SOCE), that is enhanced in amplitude and duration when SERCA is blocked using thapsigargin (Fig. 1B and 1C, TG) ^11^. This shows that SERCA-dependent ER Ca^2+^ uptake is important to limit Ca^2+^ diffusion out of the SOCE microdomain. Several studies place SERCA in close proximity to SOCE clusters ^12, 37–43^. PMCA as well has been shown to modulate Ca^2+^ levels in the SOCE microdomain in T cells ^44, 45^.

SOCE has been implicated in a broad range of physiological functions ^23^, which are likely to involve many Ca^2+^-dependent effectors. Some of these effectors localize to the SOCE microdomain and are thus activated directly by Ca^2+^ flowing through SOCE. These include calcineurin, which regulates transcription and immune cell activation through NFAT activation ^46, 47^; and adenylate cyclase 8, which coordinates crosstalk between SOCE and cAMP signaling ^48–50^. In contrast, other effectors downstream of SOCE, like Ca^2+^-activated channels at the PM do not localize to the SOCE microdomain and are often excluded from it following store depletion ^12^. For such distal effectors, Ca^2+^ tunneling contributes significantly to their activation. These Ca^2+^-activated Cl^−^ and K^+^ channels mediate fluid and ion secretion in response to SOCE ^12, 15, 51, 52^. In addition, it is important to note that the Ca^2+^ handling proteins involved in SOCE and tunneling (Orai1, STIM1, PMCA, SERCA, and IP_3_R) are themselves regulated by Ca^2+^ through modulation of their transport rate, gating, or conformation.

To better define the physiological role of Ca^2+^ tunneling, we outline its subcellular architecture and test its function in situ and in vivo. We show that SERCA localizes around the SOCE microdomain at ERPMCS defined by clustered STIM1, but is excluded from it. IP_3_R1 localizes more distally from SOCE clusters at an average distance of ∼1 µm away. PMCA is diffusely distributed at the PM and overlaps with SOCE clusters. We further generate a novel inhibitor of Ca^2+^ tunneling CaTAr by specifically targeting SERCA pumps around the SOCE microdomain. CaTAr blocks tunneling without interfering with either SOCE or IP_3_-dependent Ca^2+^ release. Blocking tunneling inhibits Cl^−^ secretion in sweat gland cells and reduces sweating in vivo in mice. Collectively, these data show that tunneling is a basic Ca^2+^ signaling modality that is important physiologically for Cl^−^ and fluid secretion.

## RESULTS

### Ca2+ tunneling in primary salivary gland cells

We had devised a protocol that temporally separates Ca^2+^ tunneling from Ca^2+^ release (Fig. 1B) ^5, 11, 12^. We showed that the Ca^2+^ tunneling signal is significantly larger and more prolonged than the signal due to SOCE alone, and that it is comparable to the Ca^2+^ signal observed following thapsigargin treatment to block ER Ca^2+^ uptake by inhibiting SERCA (Fig. 1C) ^11^. As these experiments were conducted in immortalized cell lines, we wanted to test whether tunneling is operational in a primary cell preparation. We chose acinar cells from the salivary gland as they represent a good model for Ca^2+^ tunneling given their role in saliva secretion and Ca^2+^-dependent Cl^−^ efflux ^52, 53^. Agonist mediated Ca^2+^ signals in salivary acinar cells are largely restricted to the apical membrane, where IP_3_Rs localize within a short distance (50-100 nm) from their target, the Ca^2+^-activated Cl^−^ channel ANO1 ^54, 55^. The SOCE machinery in contrast localizes predominantly to basolateral membranes ^56^. Therefore, Ca^2+^ entering through SOCE would have to navigate a considerable distance to activate the Cl^−^ channels.

Reversible inhibition of SERCA using transient CPA application increases cytosolic Ca^2+^, activates the Cl^−^ currents, and effectively depletes ER Ca^2+^ as validated by the lack of Ca^2+^ release in response to carbachol (CCh) (Supp. Fig. 1). To assess the contribution of tunneling in salivary acinar cells, we measured Ca^2+^ signals and Cl^−^ currents in response to SOCE alone versus tunneling. In both cases stores were depleted using transient CPA exposure, followed by a washout period to relieve SERCA inhibition (as per the protocol in Fig. 1B). For SOCE alone we perfused with a Ca^2+^ containing solution, whereas to induce tunneling we perfused with Ca^2+^ and CCh to activate both SOCE and IP_3_Rs. The tunneling protocol produced larger Ca^2+^ signals than SOCE alone (Fig. 1D), which were associated with greater Ca^2+^-activated Cl^−^ currents (Fig. 1E). Analyzing the kinetics of the initial rising phase of both the Ca^2+^ (Fig. 1F) and Cl^−^ signals (Fig. 1G) reveals that both develop significantly faster during the tunneling protocol compared to SOCE (7.4-fold for Ca^2+^ and 1.7-fold for Cl^−^). This shows that tunneling is faster and more efficient in activating ANO1 channels in salivary acinar cells as compared to SOCE alone.

### Architecture of the tunneling machinery

#### STIM1 and Orai1

To better define tunneling mechanistically, we were interested in mapping the architecture of the tunneling machinery. We used STIM1 and Orai1 interaction as our reference for these experiments as it has been studied extensively. STIM1 and Orai1 are diffuse throughout the ER and PM respectively at rest (Fig. 2A). Store depletion leads to their co-clustering at ERPMCS, which appear as defined puncta in Airyscan images both in the x/y (lateral) (Fig. 2A) and z (axial) dimensions (Fig. 2B). In resting cells before store depletion, the localization of STIM1 to the ER and Orai1 to the PM are apparent in orthogonal sections (Fig. 2B) and z profiles (Fig. 2C). Store depletion leads to the colocalization of STIM1 and Orai1 to ERPMCS, that is the same z plane in Airyscan images (Fig. 2B and 2D). In the super-resolution Airyscan mode the x/y resolution is estimated to be 120 nm whereas the z resolution is poorer at 350 nm ^57^. So fluorescent objects in the 350 nm Airyscan axial plane cannot be resolved. Therefore, for these experiments we used the peaks from the x/y (Fig. 2C) and z (Fig. 2D) profiles to localize the different tunneling effectors relative to each other, primarily using STIM1 as the reference for the SOCE microdomain at ERPMCS.

**Figure 2:**
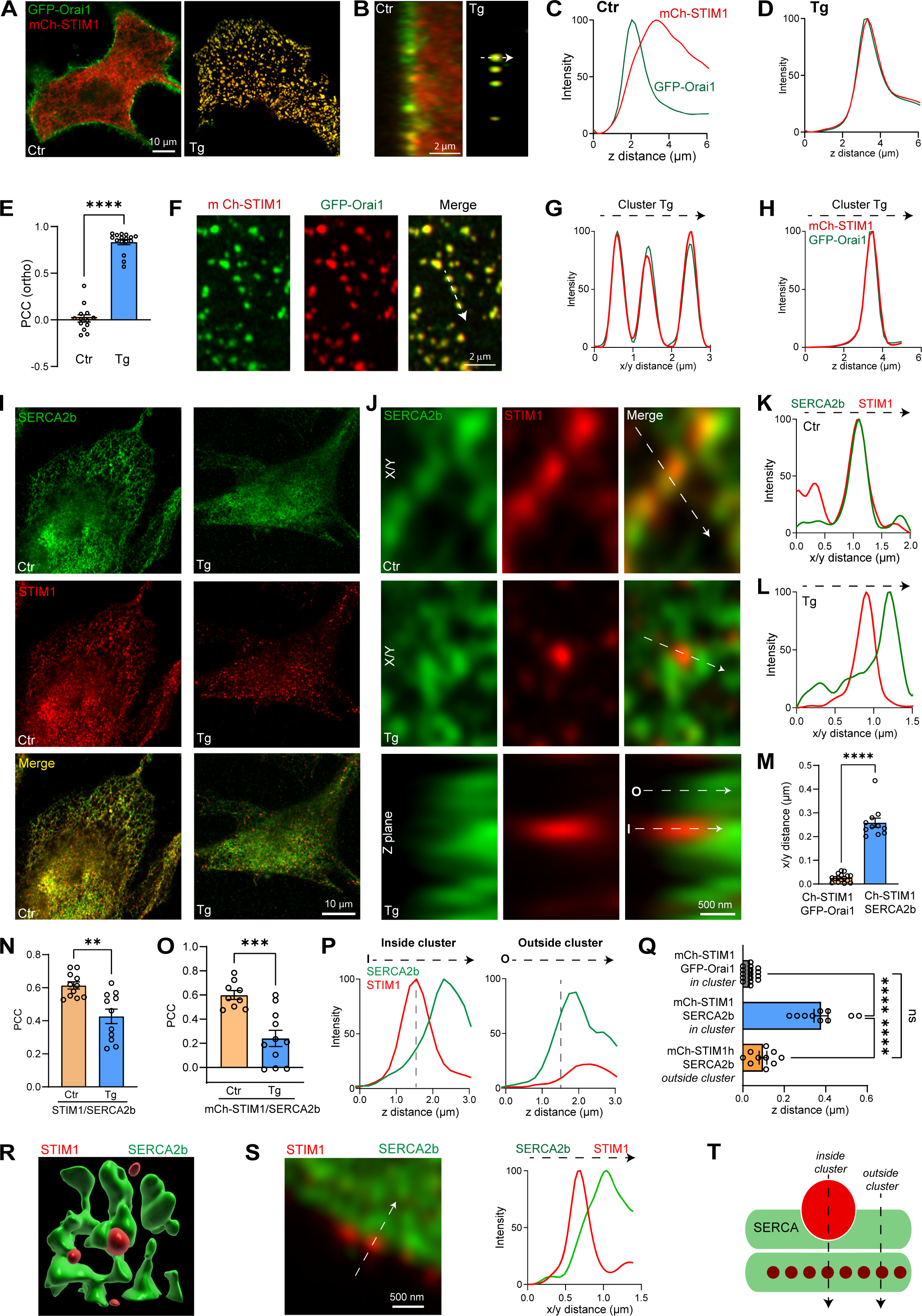
Localization of STIM1, Orai1, and SERCA2b. (**A**) Localization at the whole cell level of mCh-STIM1 and GFP-Orai1 in HeLa cells. At rest (Ctr) the confocal plane is in the middle of the cell. After store depletion with thapsigargin (Tg) the image was acquired at the PM optical plane to visualize STIM1-Orai1 co-clusters. **(B)** Orthogonal sections from confocal z-stacks in control conditions and after store depletion. **(C, D)** Relative intensities of STIM1 and Orai1 across z-stacks as in (B) before (Ctr) and after store depletion (Tg). **(E)** Pearson Correlation Coefficient (PCC) between mCh-STIM and GFP-Orai1 measured in orthogonal slices before (Ctr) and after store depletion (Tg) (n=13-16; unpaired t-test). **(F)** Enlarged confocal images showing the colocalization of STIM1 and Orai1 at the PM focal plane. **(G, H)** Plots from a virtual line scan drawn across clusters in the PM plane (G, arrow in F) and in the z axis (H, arrow in B). **(I)** Immunostaining of SERCA2b and STIM1 at the whole cell level before (Ctr) and after store depletion with Tg. **(J)** High magnification views of the spatial organization of STIM1 and SERCA2b before (Ctr) and after store depletion (Tg) in the lateral (x/y) and axial (Z plane) dimensions as indicated. **(K, L)** Intensities obtained from virtual line scans in the x/y plane across ER tubules (Ctr) and a STIM1 cluster (Tg). **(M)** Distance between the peak signals of mCh-STIM1 (center of the cluster) and the nearest SERCA2b maximum detected by immunofluorescence. The distance to the peak of GFP-Orai1 was used as a reference value (n =11-16; unpaired t-test). **(N, O)** Pearson’s Correlation Coefficient (PCC) obtained before (Ctr) and after store depletion (Tg) between endogenous SERCA2b and endogenous STIM1 (N) or overexpressed mCh-STIM1 (O). Colocalization at rest was measured in the middle of the cell and at the PM plane after store depletion (n=9-11; unpaired t-test). **(P)** STIM1 and SERCA2b normalized intensities measured over the z axis after store depletion from inside and outside a STIM1 cluster (arrows in J). **(Q)** Quantification of the peak-to-peak distance in the z axis between mCh-STIM1, Orai-GFP, and SERCA2b inside and outside the STIM1 clusters (n=13-16; one-way ANOVA). **(R)** 3D reconstruction of the relative localization of SERCA2b around the STIM1 cluster. **(S)** Image and line scan of STIM1 clusters on the side of the cell depicting its isolation from SERCA2b. **(T)** Cartoon illustrating the localization of STIM1 (red) and SERCA (green) after store depletion. The positions of the line scans in P and Q are also shown.

We calculated the Pearson colocalization coefficient (PCC) between STIM1 and Orai1 in the orthogonal plane. STIM1 and Orai1 are separate under resting conditions (PCC 0.016`0:`0.04) and significantly colocalize following store depletion (PCC 0.83`0:`0.03) (Fig. 2E). At the cluster level line scans along the membrane plane through the clusters, and in the axial dimension indicate that the position of STIM1 and Orai1 at our imaging resolution is indistinguishable in all dimensions (Figure 2F-H).

#### STIM1 and SERCA

We next localized the ER Ca^2+^ pump SERCA2b relative to the SOCE cluster by immunostaining for endogenous STIM1 and SERCA2b. Whenever possible based on the availability of validated antibodies, we localized endogenous proteins to avoid any potential artifacts due to overexpression or tagging. At rest both STIM1 and SERCA are diffusely distributed within the ER (Fig. 2I, Ctr), and colocalize as indicated in the high magnification image (Fig. 2J, Ctr) and line scan through the ER cisterna (Fig. 2K). Interestingly, store depletion separates SERCA form STIM1 in the x/y axis as observed at high magnification of a STIM1 cluster surrounded by more diffusely distributed SERCA (Fig. 2J, Tg, x/y), and through a line scan profile (Fig. 2L). Quantifying the peak to peak distance between STIM1 and SERCA in the lateral dimension confirms that SERCA, in contrast to Orai1, does not localize to the STIM1 cluster (Fig. 2M). The PCC between endogenous STIM1 and SERCA2b at the PM focal plane shows a significant reduction in their colocalization following store depletion (Fig. 2N). A similar exclusion of endogenous SERCA2b was observed from expressed mCh-STIM1 (Fig. 2O and Supp. Fig. 2).

To assess the distribution of SERCA in the axial dimension, we quantified line scans through the STIM1 cluster (inside) or in its proximity (outside) (Fig. 2J, z plane, Tg; and 2P-Q). Line scan through the STIM1 cluster shows that SERCA is excluded from the cluster and peaks deeper within the ER (Fig. 2P, left panel). Line scan just outside the STIM1 cluster shows that SERCA localizes axially with the remaining lower intensity un-clustered STIM1 deeper within the ER (Fig. 2P, right panel). These line scans were obtained from the same orthogonal z slice and normalized to the maximal intensity for STIM1 and SERCA within the slice. Comparative quantification of the axial distance between STIM1 and SERCA shows that in the axis of the STIM1 cluster (*in cluster*), SERCA localizes deeper than STIM1 as compared to colocalization of STIM1 and Orai1 in the z axis (Fig. 2Q). Outside the STIM1 cluster, the distance from the STIM1 peak to SERCA was similar to STIM1-Orai1 distance, indicating that SERCA localizes in close proximity of the PM but is excluded from the SOCE cluster (Fig. 2Q). 3D reconstruction of the STIM1 and SERCA distribution within the cortical ER supports a spatial organization where the STIM1 cluster is surrounded by SERCA (Fig. 2R). Finally, we visualized STIM1 clusters and SERCA at the edge of the cell (in the x/y plane to improve spatial resolution), which also displays clear separation between STIM1 clusters and the surrounding SERCA (Fig. 2S). Collectively these findings argue that following store depletion SERCA is not present within STIM1 clusters at ERPMCS, but is rather enriched cortically in their vicinity, both laterally and axially (Fig. 2T). Such SERCA distribution would be ideally suited to support tunneling as it would not interfere with Ca^2+^ transients within the SOCE microdomain (ERPMCS gap), but would transfer Ca^2+^ leaking out of the microdomain into the ER to fuel tunneling.

#### STIM1 and IP_3_R

Thillaiappan et al. showed that a population of cortical immobile IP_3_Rs (‘licensed’) initiate Ca^2+^ release signals and that they associate with the actin binding protein KRAP ^58, 59^. This localization as we have previously argued seamlessly supports Ca^2+^ tunneling ^3^. To assess the spatial relationship between STIM1 and IP_3_R1, we localized endogenous IP_3_R1 and KRAP relative to STIM1 clusters. At rest STIM1 and IP_3_R1 are diffuse through the ER although ‘licensed’ IP_3_R1s are visible as cortical clusters that colocalize with KRAP and appear to align along actin tracks (Supp. Fig. 3). Store depletion leads to STIM1 localization to ERPMCS without a sizable change in IP_3_R1 distribution, resulting in significant separation between STIM1 clusters and licensed IP_3_R1s (Fig. 3A). The licensed IP_3_R1 clusters colocalize perfectly with KRAP (Fig. 3B). PCC analyses confirm the colocalization of IP_3_R1 and KRAP, and the separation of STIM1 clusters from licensed IP_3_R1s or KRAP (Fig. 3C). TIRF imaging replicates the distribution of STIM1, KRAP, and IP_3_R1 relative to each other (Supp. Fig. 4A-B), supporting their cortical localization. Similar results were obtained by analyzing the colocalization of IP_3_R1-GFP with KRAP or STIM1-Ch (Supp. Fig. 4C-I). The IP_3_R1 did not colocalize with the STIM1 clusters laterally, although in the z axis licensed IP_3_R1s were in the same focal plane as STIM1, while “mobile” IP_3_Rs localized deeper in the cell (Supp. Fig. 4H-I). Of note, the expressed GFP-tagged IP_3_R1 localized closer to STIM1 clusters (Supp. Fig. 4) when compared to endogenous IP_3_R1 (Fig. 3A), arguing that either the GFP tag and/or overexpression interfere somehow with IP_3_R1 localization.

**Figure 3:**
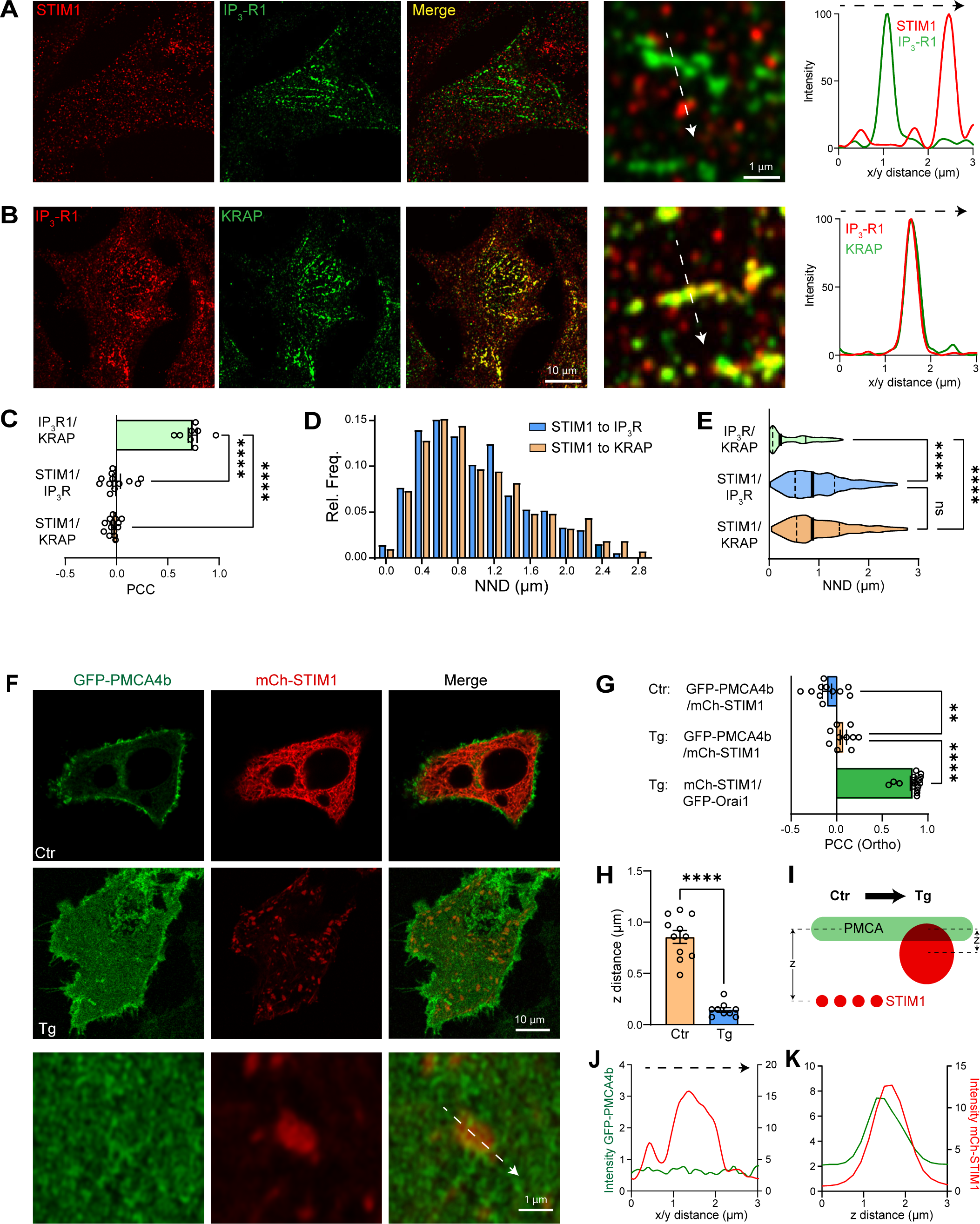
Localization of IP_3_R1 and PMCA4b. (**A**) Immunofluorescence for endogenous STIM1 (red) and IP_3_R1 (green) after store depletion with Tg. A virtual line scan (arrow) across the high magnification image shows the separation between the SOCE cluster and “licensed” IP_3_R1s. **(B)** Immunofluorescence for endogenous IP_3_R1s and KRAP at the PM focal plane. The intensity profile measured along the white arrow indicates the high degree of colocalization of the IP_3_R1 and KRAP. **(C)** PCC summaries between STIM1, IP_3_R1, and KRAP (n=8-14; one way ANOVA). **(D)** Histogram of the relative frequency of the Nearest Neighbor Distance (NND) between STIM1 clusters and IP_3_R1 or KRAP (Outliers removed using the ROUT method and a Q value of 10%). **(E)** Violin plot comparing the distribution of NNDs between the center of STIM1 clusters and IP_3_R1 or KRAP (n=398-803; one-way ANOVA). **(F)** Distribution of GFP-PMCA4b and mCh-STIM1 in HeLa cells before (Ctr) and after (Tg) store depletion. The control confocal image is taken in the middle of the cell and images after store depletion at the PM plane. **(G)** Bar chart summarizing the PCC between mCh-STIM1, GFP-Orai1 (used as a maximum reference), and GFP-PMCA4b, measured in the orthogonal plane (n=10-16; one-way ANOVA). **(H)** Quantification of the Z distance between mCh-STIM1 and GFP-PMCA4b peaks (n=9-11; unpaired t-test). **(I)** Cartoon illustrating the relative positions of PMCA and STIM1 after store depletion. **(J, K)** Intensity profiles along the x/y (J) and z (K) axis of mCh-STIM1 and GFP-PMCA4b across a STIM1 cluster (as indicated in F).

As IP_3_Rs are the site of Ca^2+^ release during Ca^2+^ tunneling the distance between IP_3_R1 and STIM1 after store depletion provides a ruler for the spatial extent of tunneling. We thus quantified the distance between STIM1 clusters and licensed IP_3_R1s (employing both IP_3_R1 or KRAP) after store depletion using Near Neighbor Distance (NND) analyses. This shows that the majority of IP_3_R1s are within 0.2-1.8 µm away from SOCE clusters (Fig. 3D), with an average distance of 0.98`0:`0.02 µm (Fig. 3E). The closest IP_3_Rs clusters where within 200 nm from the SOCE microdomain and the farthest were 2.6 µm or more away (Fig. 3D). These results argue that tunneling extends the SOCE microdomain laterally by up to 10-folds from ∼100-300 nm estimated SOCE microdomain size (ERPMCS) to a Ca^2+^ signal through IP_3_Rs up to 2.6 µm away.

#### STIM1 and PMCA

We next expressed the PM Ca^2+^ATPase 4b (GFP-PMCA4b) and measured its localization relative to STIM1-Ch before and after store depletion. At rest as expected PMCA localizes to the PM and STIM1 to the ER resulting in no colocalization (Fig. 3F-G, Ctr). Following store depletion, the PCC between STIM1 and PMCA4b increased slightly (Fig. 3G), although the absolute colocalization remained extremely low when compared with STIM1 and Orai1 colocalization (Fig. 3G). The small increase in colocalization between STIM1 and PMCA is due to the translocation of STIM1 to the PM as indicated by the decreased axial distance between the two proteins following store depletion (Fig. 3H). The intensity of the PMCA4b signal was not modified by the presence of the STIM1 cluster at the PM (Fig. 3F and 3J). In the z axis, both proteins displayed a similar profile, indicative of their close PM and ERPMCS localization (Fig. 3K). Together these analyses argue against any redistribution of PMCA4b in response to store depletion where it remains diffuse at PM. STIM1 clustering at ERPMCS in response to store depletion brings it closer to PMCA, which is diffusely distributed at the PM including within the SOCE microdomain (Fig. 3I).

### Ultrastructure of the cortical ER around ERPMCS

Functionally SERCA should localize close to SOCE cluster to allow for efficient Ca^2+^ uptake into in the ER to fuel tunneling ^5, 11, 12^. The geography of the tunneling effectors outlined above supports this conclusion and shows that SERCA surrounds SOCE clusters, without localizing to the SOCE microdomain at ERPMCS (Fig. 2I-T). We were therefore interested in obtaining a 3D view at the ultrastructural level of cortical ER including and surrounding ERPMCS where the SOCE machinery localizes. For these studies we chose a simpler cell model, Jurkats T cells, where the complexity of the cortical ER is lower, and because it is a well-studied model in terms of SOCE and has been used for careful quantifications relating to the SOCE microdomain ^35^.

We produced complete serial EM sections (50 nm thick) across three Jurkat cells and traced cortical ER, PM, and ERPMCS using the Reconstruct software (Fig. 4A) to produce 3D renditions of the cortical ER cortical around ERPMCS (Fig. 4B). We consistently observe a subdomain of the cortical ER that is linked to ERPMCS through a narrow neck and runs close to and parallel to the PM (Fig. 4B, arrows). The average depth of this ER domain relative to the PM measured from 21 junctions was 104.9`0:`11.3 nm (Fig. 4C). The shape and proximity to the PM and ERPMCS of this shallow cortical ER subdomain make it well suited for SERCA localization to position it close to but not in the SOCE microdomain at ERPMCS, although we have not directly tested this. Nonetheless, these 3D reconstructions in Jurkats reveal stratifications in the cortical ER where different tunneling effectors may localize. Whether such stratification exist in other cells remains to be determined.

**Figure 4:**
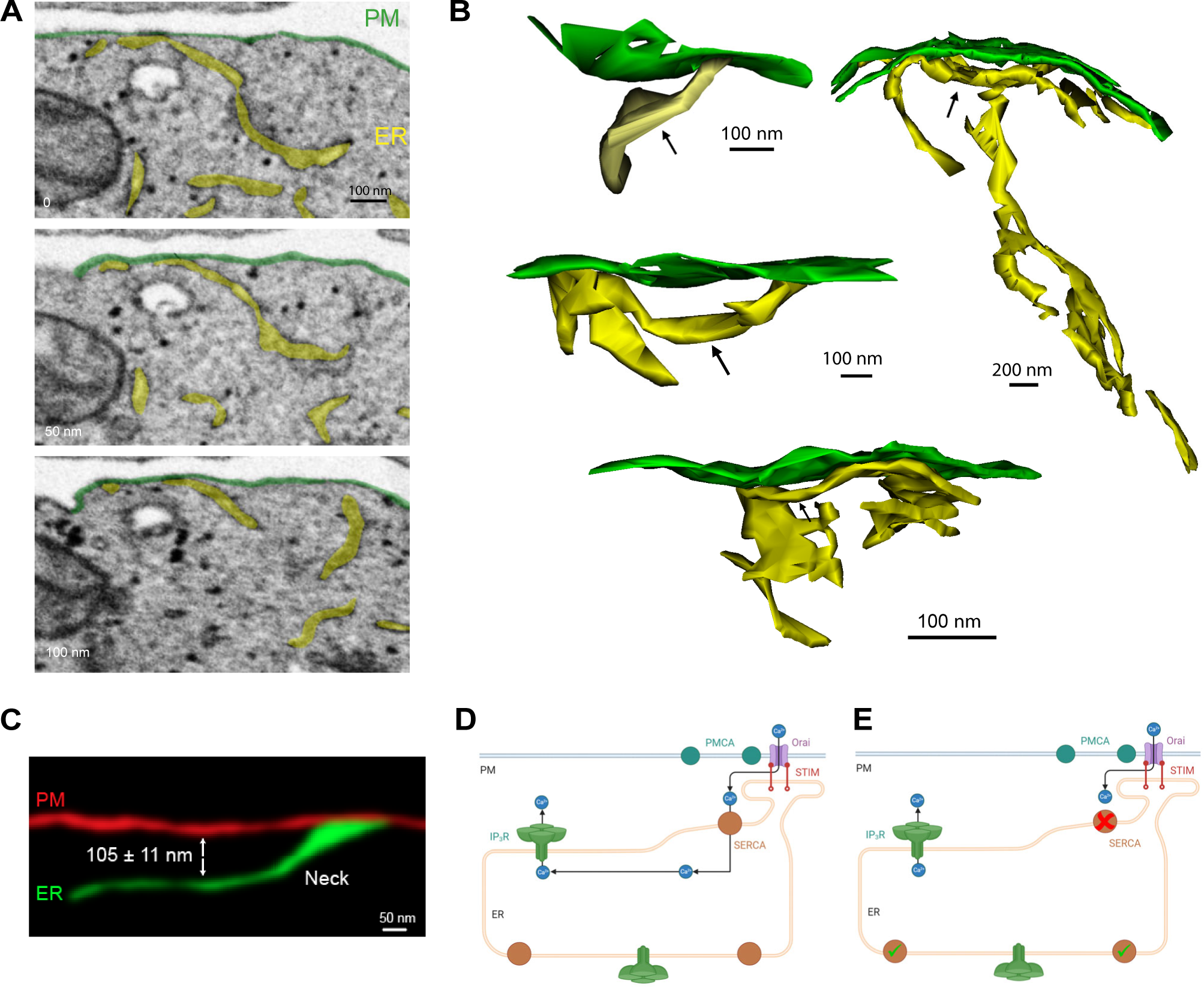
Ultrastructure 3D reconstruction of the cortical ER around ERPMCS. (**A**) Example tracings of the PM and ER used to perform 3D reconstruction. The EM images are from a sequential series across the cell. **(B)** Examples of 3D reconstructions of ERPMCS and the surrounding cortical ER (yellow) and PM (green). The arrows indicate a shallow cortical ER subdomain that typically runs parallel to the PM. **(C)** Typical example the organization an ERPMCS with the “neck” linking it to the sub-membrane cortical ER. Indicated is the average distance of the shallow cortical ER subdomain that runs parallel to the PM (n=21 junctions). **(D, E)** Cartoons illustrating the spatial organization of the tunneling machinery and the requirement to block the SERCA sub-population around the SOCE microdomain to specifically inhibit tunneling without affecting SOCE or Ca^2+^ release.

### Engineering a Ca^2+^ tunneling inhibitor (CaTAr)

Blocking Ca^2+^ tunneling specifically without affecting SOCE or Ca^2+^ release from stores is conceptually difficult as tunneling requires both pathways. It is therefore not possible to block SOCE or the IP_3_R (Fig. 4D). One could inhibit SERCA as it funnels Ca^2+^ from SOCE to distant IP_3_Rs to empower tunneling, however a global SERCA block is not viable as it would lead to store depletion and constitutive SOCE activation due to the inherent ER Ca^2+^ leak. But, would it be possible to inhibit only the SERCA pool that surrounds the SOCE clusters at ERPMCS? Should this be possible it would produce a specific tunneling blocker without affecting SOCE or Ca^2+^ release (Fig. 4E).

We noticed from studies co-localizing the artificial ER-PM junction marker MAPPER and STIM1, that surprisingly the two proteins do not colocalize after store depletion. At rest when stores are full MAPPER localizes to ERPMCS with some faint un-clustered STIM1 at high magnification (Fig. 5A, Ctr). Interestingly, following store depletion MAPPER and STIM1 localize to distinct subdomains laterally within ERPMCS (Fig. 5A, CPA), where MAPPER is clearly excluded from STIM1 clusters (Fig. 5A), as previously reported ^31, 34^. Time lapse imaging of the evolution of STIM1 clustering at ERPMCS show that STIM1 moving along microtubule tracks at rest, clusters in response to store depletion within the ERPMCS and pushes out MAPPER to form a subdomain devoid to MAPPER within the same ERPMCS (Supp. Video 1). Orthogonal sections through the clusters confirm that after store depletion STIM1 and MAPPER are within the same axial plane but do not co-localize (Figure 5B). We measured the STIM1 to MAPPER edge-edge distance which was close to 0 while STIM1-STIM1 and MAPPER-MAPPER were far apart (Supp. Fig. 5A). This argues that STIM1 clusters within a MAPPER filled contact site.

**Figure 5:**
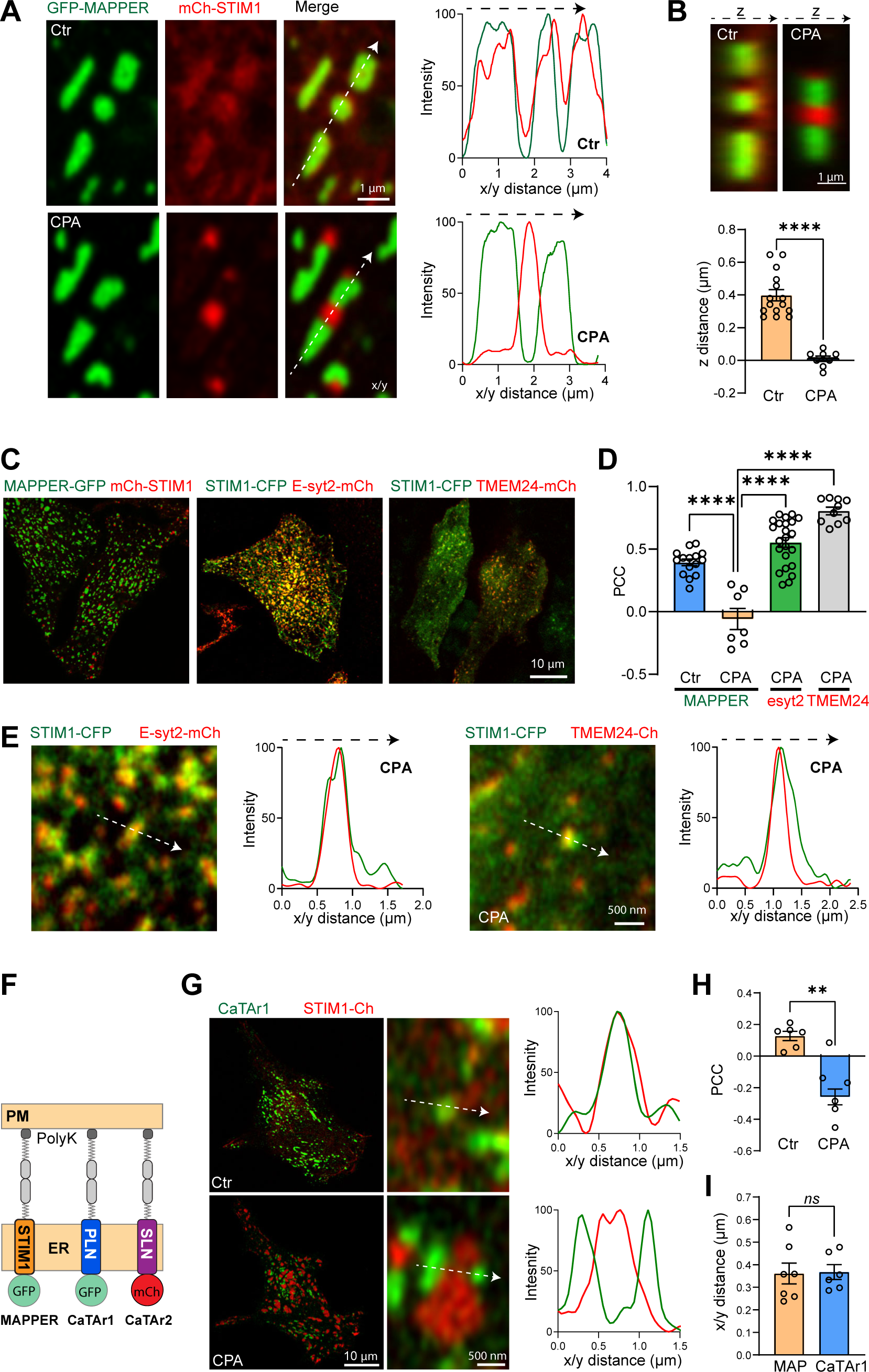
Designing a tunneling specific blocker CaTAr. (**A**) High magnification Airyscan images of GFP-MAPPER and mCh-STIM1 at rest (Ctr) and after store depletion (CPA). Line scans as indicated on the merged image show colocalization of MAPPER (green) with the diffuse un-clustered STIM1 (red) at rest, and the separation of clustered STIM1 from MAPPER. **(B)** Orthogonal sections of GFP-MAPPER and mCh-STIM1 before (Ctr) and after store depletion (CPA). The bar chart reports the distance in the Z axis between maximum intensities of GFP-MAPPER and mCh-STIM1 signals before (Ctr) and after store depletion (CPA) (n=8-15; unpaired t-test). **(C)** Localization of MAPPER-GFP, E-Syt2-mCh, and TMEM24-mCh at the whole cell level. **(D)** Bar chart summarizing the PCC of the three tethers relative to STIM1 (n=7-23; one-way ANOVA). **(E)** Confocal images and line scans illustrating the colocalization of E-Syt2-mCh and TMEM24-mCh with STIM1-CFP clusters. The intensity plots are obtained from the lines depicted by the white arrows. **(F)** Cartoon depicting the structure of the Ca^2+^ Tunneling Attenuators (CaTAr1 and 2) compared to MAPPER. **(G)** Confocal images at the PM plane of CaTAr1 and mCh-STIM1 before (Ctr) and after (CPA) store depletion. The intensity plots using the white arrows in the high magnification images report the colocalization of STIM1 and CaTAr1 at rest and their separation after store depletion. **(H)** Bar chart summarizing the PCC between CaTAr1 and mCh-STIM1 before (Ctr) and after store depletion (CPA) (n=6; paired t-test). **(I)** Bar chart summarizing the distance between STIM1 clusters and CaTAr1 or MAPPER (MAP) (n=6-7; unpaired t-test).

We obtained a similar separation between STIM1 clusters and a shorter version of MAPPER (MAPPER-S) (Supp. Fig. 5B-C), that brings the ER and PM closer together (within 10 nm) ^60^ as previously shown ^31^, arguing that the separation between MAPPER and STIM1 is not dependent on the gap distance between the ER and PM.

The distribution of MAPPER relative to STIM1 within ERPMCS argues that STIM1-Orai1 clusters may form a privileged subdomain within an ERPMCS that prevents other molecules from colocalizing with it. To test whether this is the case, we assessed the localization of STIM1 relative to two endogenous ER-PM tethers, E-Syt2 and TMEM24 ^33, 60^. In contrast to MAPPER, both E-Syt2 and TMEM24 co-localize with STIM1 clusters within ERPMCS as assessed at the whole cell level (Fig. 5C), at high magnification within STIM1 clusters (Fig. 5E), using line scan across individual clusters (Fig. 5E), and by quantifying colocalization using PCC (Fig. 5D).

The distinct distribution of MAPPER following store depletion raised the intriguing possibility that it could localize close to or within SERCA rich cortical ER subdomains, especially that MAPPER has been shown to extend ERPMCS ^31, 34^. Should this be the case it would provide an ideally targeted molecule to inhibit cortical SERCA. But how to specifically inhibit SERCA using MAPPER? For this we used the transmembrane domain of two well characterized SERCA inhibitors in muscle cells Phospholamban (PLN) and Sarcolipin (SLN), that importantly modulate SERCA function through their transmembrane domains (TM) ^61^. Therefore, to create a potential cortical SERCA inhibitor-that is a specific tunneling blocker-, we replaced the MAPPER TM domain with either the TM domain of PLN or SLN (Fig. 5F). This generated GFP-PLN-MAPPER, that we named CaTAr1 for <u>Ca</u>^2+^ <u>T</u>unneling <u>A</u>ttenuator <u>1</u>; and Ch-SLN-MAPPER named CaTAr2. When co-expressed with mCh-STIM1 or STIM1-CFP both CatAr1 (Fig. 5G) and CaTAr2 (Supp. Fig. 6) had the same localization relative to clustered STIM1 as MAPPER. They segregate apart from STIM1 clusters following store depletion (Fig. 5G-I, Supp. Fig. 6A, Supp. Video 2). CaTAr2 also colocalizes with MAPPER (Supp. Fig. 6B) and was isolated from cortical IP_3_R1-GFP (Supp. Fig. 6C). Together those results argue that the CaTAr constructs localize around but not within STIM1 cluster in ERPMCS.

To confirm that PLN inhibits SERCA in HeLa cells, we expressed wild-type PLN and show that it distributes throughout the entire ER, where it colocalizes with STIM1 (Supp. Fig. 7A), and leads to store depletion with a significant reduction of histamine induced Ca^2+^ release in PLN expressing cells (Supp. Fig. 7B-D), consistent with global SERCA inhibition.

### CaTAr inhibits Ca^2+^ tunneling

We next asked whether the engineered CaTAr constructs block Ca^2+^ tunneling and whether they affect SOCE or Ca^2+^ release. We first tested whether the CaTAr backbone, i.e. MAPPER, affects tunneling. Applying our standard tunneling protocol to HeLa cells expressing MAPPER-GFP and loaded with the Ca^2+^ indicator Calbryte590, leads to similar Ca^2+^ tunneling amplitudes in control and MAPPER expressing cells (Fig. 6A-C), showing that MAPPER expression does not significantly affect Ca^2+^ tunneling.

**Figure 6:**
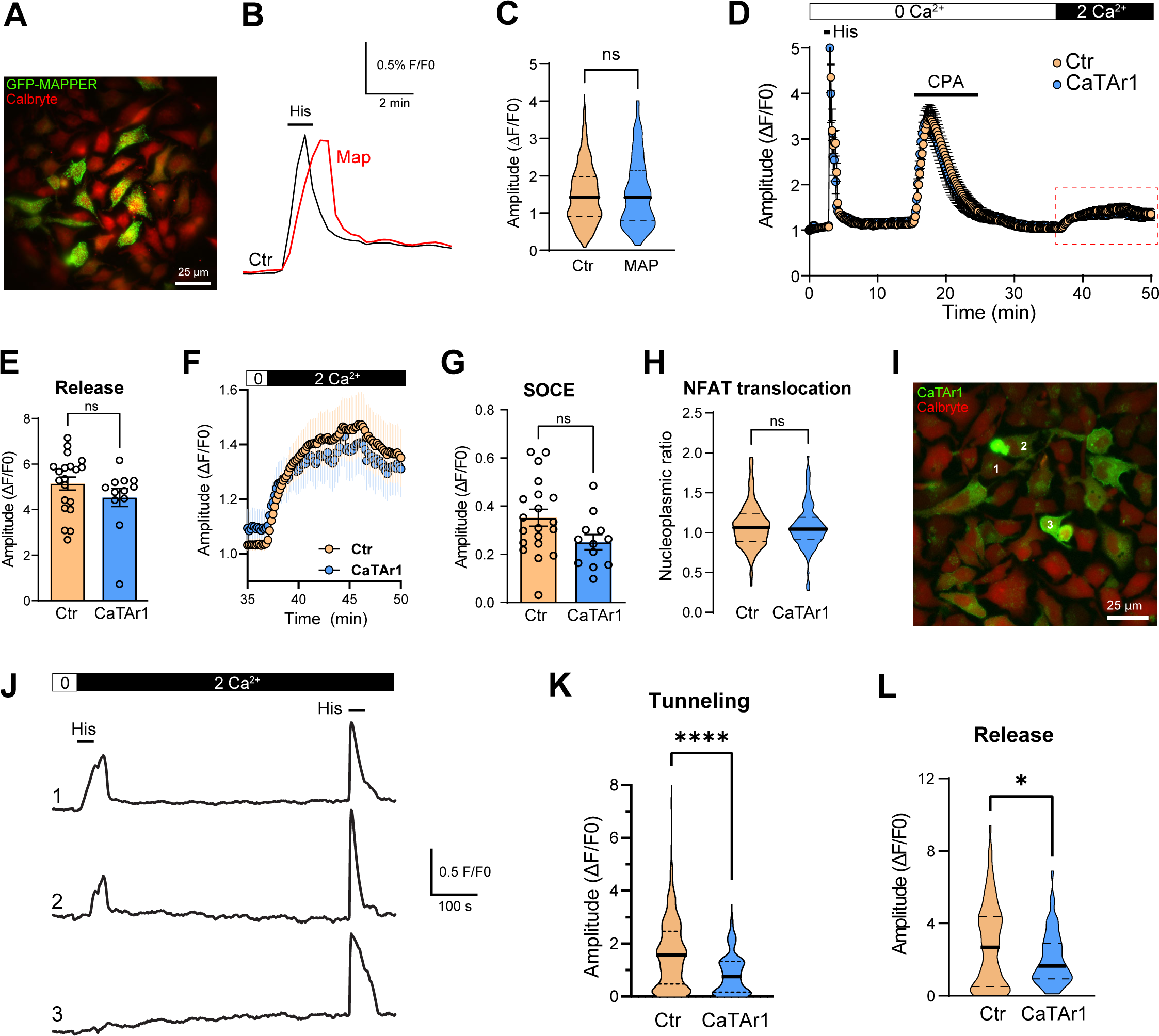
CaTAr1 inhibits Ca^2+^ tunneling. (**A**) HeLa cells expressing GFP-MAPPER and loaded with the Ca^2+^ indicator Calbryte 590. **(B)** Changes in intracellular Ca^2+^ during a tunneling experiment from untransfected cells (Ctr) and cells expressing GFP-MAPPER (MAP). **(C)** Violin plot of the amplitude of the tunneling signal in Ctr and MAPPER expressing cells (MAP) (n=118-439; unpaired t-test). **(D)** Ca^2+^ release and SOCE in control (Ctr; no detectable CaTAr1 expression) and CatAr1 expressing cells from the same dish. Histamine-induced (His, 100 μM) Ca^2+^ release from stores was followed by CPA to deplete Ca^2+^ store, a wash period in Ca^2+^ free media to allow for SERCA to be active, and then the addition of Ca^2+^ to activate SOCE (n=5-9). **(E)** Bar chart summarizing the levels of Ca^2+^ release in response to histamine in a Ca^2+^-free solution (n=12-20; unpaired t-test). **(F, G)** Enlarged traces from the red rectangle in (D) and bar chart summarizing the levels of SOCE in cells that did not express CatAr1 (Ctr) and CaTAr1 expressing cells (n=12-20; unpaired t-test). **(H)** Quantification of NFAT nuclear translocation in Ctr and CaTAr1 expressing cells (n=109-269; unpaired t-test). **(I)** HeLa cells expressing CaTAr1 and loaded with Calbryte 590. The numbers correspond to the cells in (J). **(J)** Traces showing the Ca^2+^ tunneling transient in response to His+Ca^2+^ following the standard store depletion with CPA and wash (not shown for clarity). A second histamine application after a delay in Ca^2+^-containing media confirms that all cells including those expressing CatAr1 refill their stores. **(K, L)** Violin plots summarizing the levels of Ca^2+^ tunneling (K) and Ca^2+^ release in response to His following the second application in Ca^2+^-containing media (L) (n=109-269 (K); n=56-182 (L); unpaired t-test).

We then tested the effect of CaTAr1 expression on agonist dependent Ca^2+^ release and SOCE. CaTAr1 expression did not alter the levels of Ca^2+^ release in response to histamine in Ca^2+^-free media (Fig. 6D-E), showing that it does not modulate agonist dependent Ca^2+^ release. Similarly, CPA dependent Ca^2+^ release was not affected by CaTAr1 expression arguing against any modulation of ER Ca^2+^ leak or Ca^2+^ store content (Fig. 6D). We then measured SOCE after store depletion with CPA and a wash period to allow SERCA to be active. SOCE levels were not affected by CaTAr1 expression (Fig. 6F-G). However, as this SOCE signal was quite small, modest effects on SOCE due to CaTAr1 expression may be difficult to quantify. To support our conclusion that CaTAr1 expression does not affect SOCE, we used NFAT nuclear translocation as a functional reporter of SOCE levels within ERPMCS. CaTAr1 expression did not affect NFAT nuclear translocation (Fig. 6H).

Ca^2+^ tunneling in contrast was significantly and dose-dependently inhibited by CaTAr1 expression (Fig. 6I-K). At the individual cell level, the extent of inhibition of Ca^2+^ tunneling correlates with the expression levels of CaTAr1 (Fig. 6I-J), with complete inhibition in cells with high CaTAr1 expression (Fig. 6I-J, cell#3). At the population level, the amplitude of the tunneling signal was significantly reduced (by ∼47 %) in cells expressing CaTAr1 compared to those with no expression in the same dish (Fig. 6K).

A second application of histamine following Ca^2+^ tunneling confirms the ability of the CatAr1 expressing cells to refill their stores (Fig. 6J). This release after tunneling and store refilling shows a small reduction in amplitude in cells expressing CaTAr1 (Fig. 6L). Interestingly, as discussed above, His-induced Ca^2+^ release in Ca^2+^-free media had a similar amplitude in control and CatAr1 expressing cells (Fig. 6D-E). In Ca^2+^-free media the Ca^2+^ signal depends solely on Ca^2+^ release from stores as there is no influx. As we observe a decrease in the Ca^2+^ release signal in response to histamine in Ca^2+^-containing but not Ca^2+^-free media, indicates that a fraction of the Ca^2+^ during the release phase in response to agonist is through tunneling. This is important as it argues that tunneling is activated early on following Ca^2+^ release due to partial store depletion while IP_3_ levels remain high.

Similar results were obtained with CaTAr2, which inhibited Ca^2+^ tunneling with no effect on Ca^2+^ release (Supp. Fig. 6D-F). CaTAr2 was less potent than CaTAr1 with ∼24 % reduction in Ca^2+^ tunneling in CaTAr2 expressing cells as compared to non-expressing cells in the same dish (Supp. Fig. 6E).

Collectively, these results show that the CaTAr constructs are specific tunneling inhibitors that reduce Ca^2+^ tunneling without affecting either Ca^2+^ release from stores or SOCE. This provides a powerful tool to test the role and contribution of tunneling to cellular and physiological responses.

### CaTAr inhibits Cl^−^ secretion in human sweat cells

Sweating depends on SOCE as patients with reduced SOCE due to mutations in either STIM1 or Orai1 suffer from anhidrosis ^24^. The contributions of SOCE and the Ca^2+^-activated Cl channel ANO1 to sweat production has been demonstrated in situ in mice using genetic or chemical inhibition of SOCE, and in the immortalized human eccrine sweat gland cell line NCL-SG3, respectively ^52, 62^. Therefore, NCL cells represent a good model to test the role of tunneling on sweating and the effectiveness of CaTAr. We previously showed that ANO1 segregates away from SOCE clusters in response to store depletion in *Xenopus* oocytes, thus requiring tunneling to deliver Ca^2+^ entering through SOCE to ANO1 ^12^.

We localized ANO1 relative to STIM1 in NCL cells using either an antibody against ANO1 or by expressing ANO1-GFP together with Ch-STIM1. As expected, ANO1 localizes to the PM and STIM1 to the ER at rest (Fig. 7A and Supp. Fig. 8A). Following store depletion ANO1 was largely excluded from SOCE clusters and localized at their periphery (Fig. 7A and Supp. Fig. 8A). We quantified ANO1-STIM1 colocalization using PCC as compared to STIM1-Orai1. Orai1 colocalizes with STIM1 in NCL cells to a similar level as in HeLa cells (Fig. 7B). In contrast, ANO1 (endogenous or GFP-tagged) was largely isolated from SOCE clusters (Fig. 7B). This separation is further illustrated by the STIM1 to ANO1 peak-to-peak distance in the lateral dimension (x/y) (Fig. 7C), while in the axial dimension STIM1, Orai1, and ANO1 localize to the same Airyscan focal plane (Fig. 7D).

**Figure 7:**
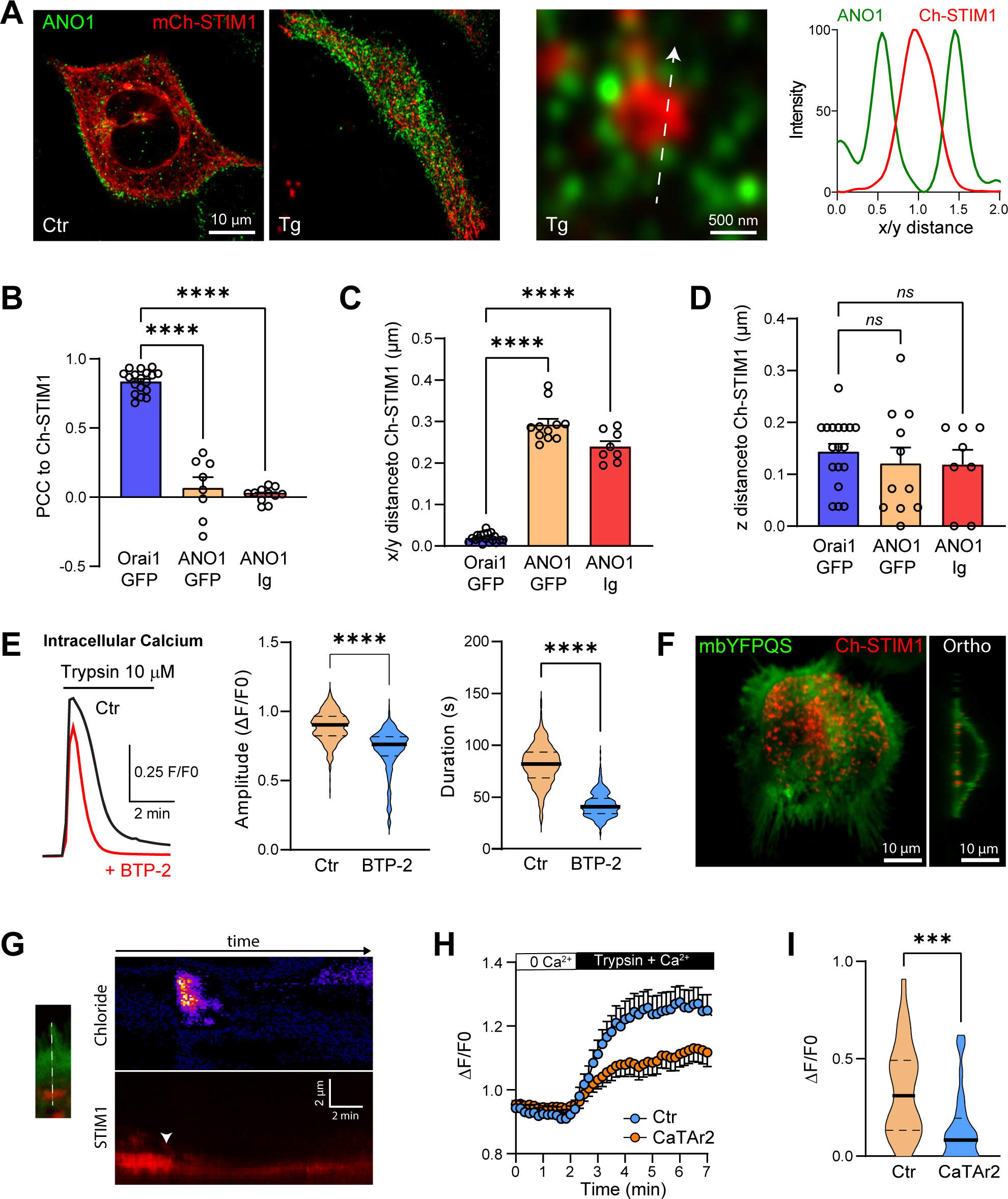
Tunneling in NCL-SG3 cells. (**A**) Localization of ANO1 by immunocytochemistry and mCh-STIM1 before (Ctr) and after store depletion (Tg). An intensity plot performed along a line (arrow) crossing a STIM1 cluster illustrates the separation of STIM1 and ANO1 at the PM focal plane. **(B-D)** Comparative localization at the PM plane of mCh-STIM1 with ANO1-GFP and the endogenous ANO1 protein (ANO1-Ig) following store depletion. As indicated by the PCC and the peak-to-peak distances the two proteins do not colocalize but are localized at the same optical plane (z distance) (n=8-18; one-way ANOVA). **(E)** Intracellular Ca^2+^ elevation induced by trypsin application in NCL-SG3 cells loaded with the Ca^2+^ probe Fluo4-AM. The application of the SOCE inhibitor BTP-2 (10 μM) reduces the amplitude and the duration of Ca^2+^ release (n=379-402; unpaired t-test). **(F)** Confocal images of NCL-SG3 cells expressing the Cl^−^ sensor mbYFPQS and mCh-STIM1 after store depletion. The orthogonal section through the cell indicates the PM localization of the chloride sensor. **(G)** Kymographs measured during a tunneling event on cells expressing mbYFPQS and mCh-STIM1. The line passes through a SOCE cluster and an adjacent cell appendage labelled by mbYFPQS. The changes in Cl^−^ concentration (upper) are distal from the STIM1 cluster, which indicates the Ca^2+^ entry point. **(H)** Time course of the amplitude of the Cl^−^ signal induced by Ca^2+^ tunneling from NCL-SG3 cells in the same dish that do not show any CaTAr2 expression (Ctr) and cells expressing CaTAr2. **(I)** Violin plots summarizing the amplitude of the Cl^−^ signal 3 min after trypsin and Ca^2+^ addition to stimulate tunneling (n=32-33; unpaired t-test).

To induce Cl^−^ secretion in NCL cells we used trypsin to activate the Proteinase Activated Receptor 2 (PAR2) which leads to Ca^2+^ release (Fig. 7E). Trypsin induces a rise in the cytosolic Ca^2+^ that was reduced in amplitude (18.8±0.9%), and in a more pronounced fashion in its duration (48±0.8%) by the CRAC channel inhibitor BTP2 (Fig. 7E). This argues for a role for SOCE/tunneling in generating the maximal Ca^2+^ signal in response to agonist stimulation. We then expressed in NCL cells the membrane bound Cl^−^ sensor mbYFPQS ^63^, which localizes diffusely to the PM, including to the microvilli away from the STIM1 clusters (Fig. 7F). Orthogonal sections through confocal z-stacks confirmed the membrane localization of the sensor (Fig. 7F). We subjected cells co-expressing mbYFPQS and STIM1 to our standard tunneling protocol and used a line scan across an SOCE cluster and the cell margin to follow the temporal progression of Cl secretion using TIRF microscopy (Fig. 7G). The kymographs show a rise in mbYFPQS fluorescence (indicative of Cl secretion) far from the STIM1 cluster (the site of Ca^2+^ entry) (Fig. 7G). This supports Ca^2+^ tunneling from the SOCE point source entry to the distal ANO1 channels (Fig. 7G). Note that the intensity of the STIM1 cluster decreases along the kymograph following store refilling (Fig. 7G), which is indicative of clustered STIM1 dissociation. These data show that Ca^2+^ tunneling supports Cl secretion distally to the SOCE clusters.

To assess the contribution of Ca^2+^ tunneling to Cl^−^ secretion we used CaTAr2 (mCh-SLN-MAPPER), to avoid overlap with the mbYFPQS fluorescence. Expression of CaTAr2 led a slower and smaller Cl^−^ secretion signal (Fig. 7H). Quantification of Cl^−^ secretion from mbYFPQS fluorescence at 3 min post-trypsin shows a significant decrease in CaTAr2 expressing cells (Fig. 7I). Collectively these results show that tunneling is an important contributor to the activation the Ca^2+^-activated Cl^−^ channels in NCL sweat gland cells.

### Tunneling supports sweating in vivo

We next wanted to evaluate the potential contribution of tunneling in vivo. We used the paw sweat test as a model since it has been successfully used in the past as an indicator of SOCE’s contribution to sweating in mice ^52, 64^. We used an adenoviral vector to express CaTAr1 under the control of the CMV promoter in the paw of mice. CaTAr1 expression (GFP) could be detected 4 days following the injection of the adenovirus (Fig. 8A). We injected an adenovirus expressing GFP alone as a control. Sweating was evaluated by applying starch iodine to the paw and measuring the area covered by the black dots (Fig. 8B-C). We observe a significant reduction in sweating in animals expressing CaTAr1 as compared to GFP (Fig. 8B-D). This supports the reduction in Cl^−^ secretion following tunneling inhibition in NCL cells and shows that tunneling is an important contributor to sweat production in vivo.

**Figure 8:**
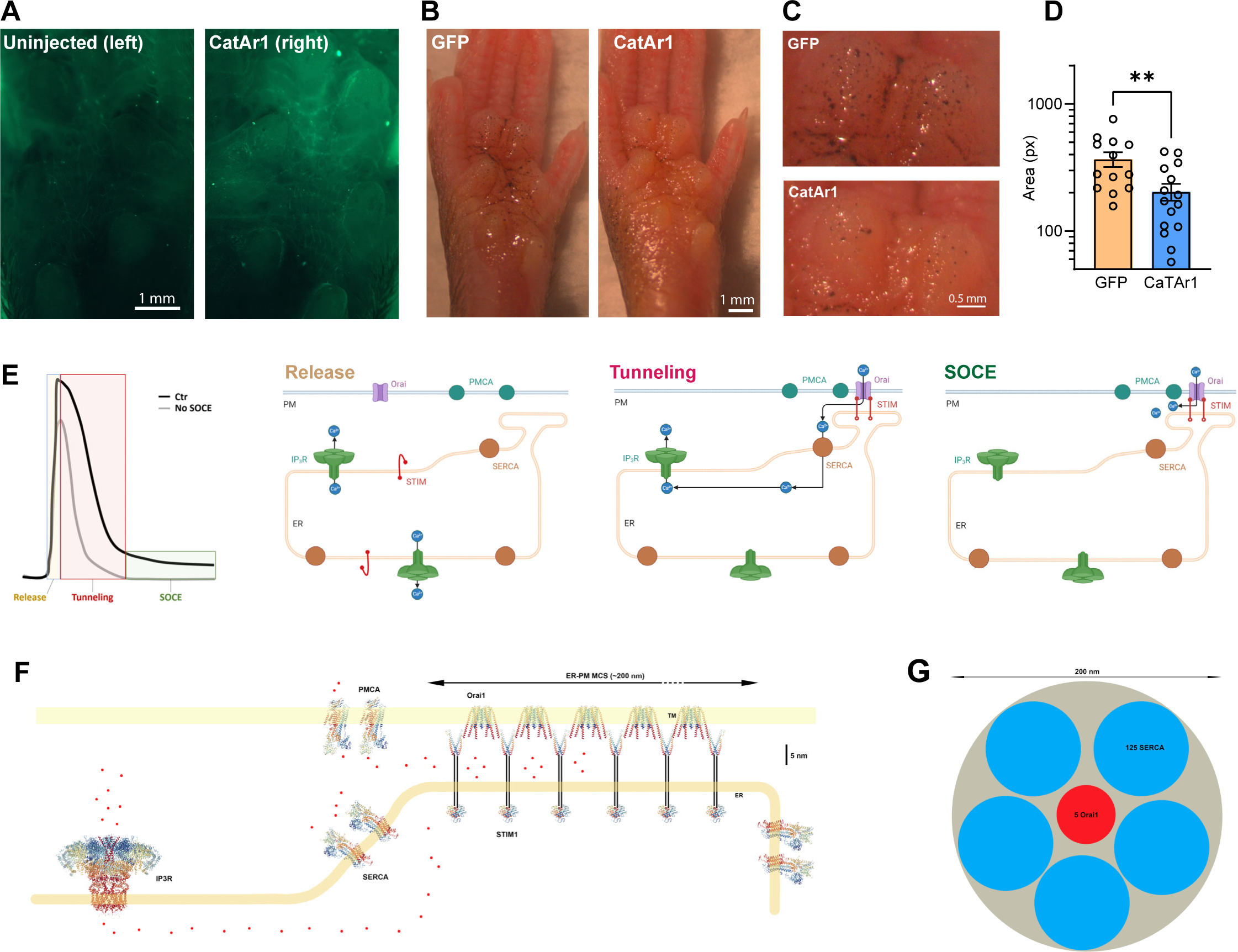
Inhibition of sweating by CaTAr1. (**A**) GFP fluorescence recorded in the mouse paw injected with CaTAr1 (right) compared with the contralateral paw (left) on the same animal. **(B, C)** Sweating visualized using the iodine/starch technique in GFP– and in CatAr1-expressing paws. **(D)** Bar chart summarizing the sweating areas recorded after 15 min in control (GFP) and CaTAr1 expressing paws (n=13-15; unpaired t-test). **(E)** Cartoons illustrating the different phases of a typical agonist driven Ca^2+^ release signal with (Ctr) or without SOCE. The activation state and localization of the Ca^2+^ effectors are shown for the different phases. **(F)** To scale illustration of the Ca^2+^ effectors involved in tunneling. For STIM1 only the domains with the atomic structure solved are shown. **(G)** To scale depiction of 5 Orai1 channels and clusters of 125 SERCAs each.

## Discussion

Various agonists produce intracellular Ca^2+^ signals through activation of G-protein or receptor tyrosine kinase coupled receptors that induce PLC to produce IP_3_ and releases Ca^2+^ from ER Ca^2+^ stores. The release phase is followed by Ca^2+^ influx from the extracellular space through activation of SOCE in response to store depletion. PLC generated lipid messengers can also activate members of the TRP family some of which are Ca^2+^ permeant ^65, 66^. However, SOCE is highly Ca^2+^ selective and is responsible to the prolonged low amplitude Ca^2+^ signal following store depletion. The SOCE signal inactivates when IP_3_ levels drop, leading to termination of Ca^2+^ release through IP_3_Rs, store refilling by SERCA, and dissociation of the STIM1-Orai1 clusters at ERPMCS. The work presented here builds on previous studies and shows that agonist mediated Ca^2+^ signaling has a third central component, Ca^2+^ tunneling (Fig. 8E).

We have previously shown that Ca^2+^ tunneling is primarily cortical and have argued that this is because of the large IP_3_R conductance for Ca^2+^ compared to that of Orai1, and because of the distribution of IP_3_Rs ^3, 11^. The geography of the primary Ca^2+^ tunneling effectors (SERCA, PMCA, IP_3_R1) relative to STIM1 outlined herein supports and extends this conclusion. We localized endogenous STIM1, SERCA, and IP_3_R1 to avoid issues with overexpressed tagged proteins. We show that a population of ‘licensed’ IP_3_R1 colocalize with KRAP cortically close the PM, consistent with previous studies ^3, 58, 59^. Near neighbor distance (NND) analyses show that on average licensed IP_3_R1s localize ∼1 µm away from a STIM1 cluster and can be up to over 2 µm away. This effectively expands the ability of Ca^2+^ flowing through SOCE to reach cortical effectors that are 1-2 µm away from an SOCE cluster at ERPMCS. Ca^2+^ that enters the cell within the SOCE microdomain and is taken up into the ER lumen by SERCA is released by the nearest open IP_3_R1 it encounters. Given the large conductance of IP_3_R1 the limiting factors for this Ca^2+^ tunneling would be the rate of Ca^2+^ entry through Orai1 and SERCA uptake. Comparatively, the Ca^2+^ signal due to SOCE alone would be limited to the SOCE microdomain (100-300 nm in diameter) and thus would activate effectors within the SOCE microdomain or in their immediate vicinity. Therefore, the localization of endogenous IP_3_R1 cortically close the PM axially but distal to the SOCE microdomain laterally, is well suited to support tunneling by expanding the spatial extent of SOCE (up to 10-fold) to reach distal Ca^2+^ dependent effectors. We show that this is indeed the case as the Ca^2+^ tunneling protocol activates larger and faster Ca^2+^ signals and associated Cl^−^ currents in primary salivary cells (Fig. 1D-G). We further show that blocking tunneling in sweat cells (Fig. 7H-I) or in vivo in mice (Fig. 8A-D) inhibits Cl^−^ secretion and sweating, respectively.

Previous imaging studies in different cell types, including our own study, used tagged overexpressed SERCA and argued for close association between SERCA and STIM1 placing SERCA within or surrounding the SOCE microdomain ^12, 37, 39–41^. In some studies, this was supplemented by FRET and co-IP experiments as well ^37, 42^. We were concerned that overexpression or the tag itself could interfere with localization especially when using high resolution microscopy, as we observed a differential localization of tagged IP_3_R1 (Supp. Fig. 4) as compared to endogenous IP_3_R1 (Fig. 3A-E). Tagged IP_3_R1 localized closer to STIM1 clusters compared to endogenous IP_3_R1. In contrast, we did not observe a differential localization between tagged and endogenous STIM1. We therefore opted to localize endogenous SERCA2b as good antibodies are available. We show that SERCA2b is excluded from the SOCE microdomain within ERPMCS, and rather surrounds it within a cortical ER domain that localizes close the ERPMCS. This is consistent with functional studies arguing for SERCA being close to the SOCE microdomain ^11, 38^. The localization of SERCA suggests a funnel around the SOCE microdomain that collects Ca^2+^ ions that spillover out of ERPMCS and tunnel them into the ER to support IP_3_R–mediate release at distal sites. Such a distribution would fit well with the slow transport rate for SERCA (∼40 Ca^2+^/sec at V_max_ ^35, 67^) requiring around 125 SERCA molecules to take up the Ca^2+^ flowing through a single Orai1 channel (∼5,000 ions/sec ^35^). As SERCA localizes outside the SOCE microdomain it would be expected to transport only a fraction of Ca^2+^ ions entering the cell through Orai1 channels; that is the fraction that spills out of the SOCE microdomain. Is it possible to fit such a large number of SERCAs around a STIM1-Orai1 cluster? Figure 8F shows a to scale rendition of the different tunneling effectors based on published structures. Assuming 5 Orai1 channels within the SOCE microdomain and 125 SERCA/Orai1 channel the SERCAs required would fit within a 200 nm diameter ER domain (Fig. 8G).

The gap between the ER and PM has been shown using cryo-EM tomography to be modulated by different E-Syt isoforms ^29^. When the ERPMCS gap was artificially shorten using chemically induced linkers to below 6 nm, this prevented stabilization of the STIM1-Orai1 clusters within the junction and favored their enrichment at the junction periphery ^32^. Furthermore, similar to our findings Henry et al. showed that MAPPER localizes to a different subdomain than STIM1 within ERPMCS, whereas a longer tether Sec22 colocalized with STIM1 in ERPMCS ^31^, similar to what we show herein for two endogenous tethers TMEM24 and E-Syt2 (Fig. 5C-E). Based on these findings Henry et al. argued that the STIM1-Orai1 clusters localize to the periphery of ERPMCS. However, our findings regarding the localization of SERCA and that of CaTAr (as an inhibitor of cortical junctional SERCAs) suggest that the STIM1-Orai1 clusters localize to the center of ERPMCS with the SERCA-CaTAr complex being more peripheral. We took advantage of this unusual MAPPER localization at the periphery of STIM1 clusters to develop a specific tunneling inhibitor (CaTAr). We replaced the MAPPER TM domain with either PLN of SLN to specifically block the subpopulation of cortical SERCAs close to the SOCE microdomain. This effectively reduced tunneling without affecting Ca^2+^ release from stores or SOCE (Fig. 6).

We used CaTAr to validate the functional importance of Ca^2+^ tunneling in sweat production. Blocking tunneling reduces Cl^−^ secretion in NCL sweat cells (Fig. 7H-I), and more importantly, it reduces sweating in vivo in mice injected with a virus expressing CaTAr in the paw (Fig. 8A-D). Furthermore, we show that a tunneling protocol activates larger Cl^−^ currents in primary salivary cells (Fig. 1). This is consistent with an early study on HSG cells from the submandibular salivary gland showing that ER Ca^2+^ and IP_3_-dependent Ca^2+^ release are needed for optimal activation of Ca^2+^-activated K^+^ currents ^51^, a mechanism similar to Ca^2+^ tunneling. In addition, Ca^2+^ tunneling has been implicated in secretion in the exocrine pancreas ^4, 13, 68^. Collectively, these findings argue for an important role for Ca^2+^ tunneling in fluid secretion in exocrine glands by modulating the activity of Cl^−^ and K^+^ channels to support vectorial ion and fluid secretion.

Based on both the architecture of the Ca^2+^ signaling machinery following store depletion and the functional data following tunneling inhibition, we propose three phases in the Ca^2+^ signal following agonist stimulation (Fig. 8E). At rest with Ca^2+^ stores full, IP_3_ production gates IP_3_Rs and release Ca^2+^ from stores resulting in the initial rise of the cytosolic Ca^2+^ signal (Fig. 8E, Release). This leads to gradual store depletion and activation of SOCE. During the early phases of store depletion, IP_3_R would still be open due to the presence of IP_3_ from receptor activation. These conditions would support Ca^2+^ tunneling as SOCE is active and IP_3_Rs are open (Fig. 8E, Tunneling). When IP_3_ levels in the cytosol fall below the threshold to open IP_3_Rs, this leads to the termination of tunneling and maintenance of only SOCE with a lower amplitude global Ca^2+^ signal as Ca^2+^ entry would localize primarily to the SOCE microdomain (Fig. 8E, SOCE). Functional evidence supports this model. In salivary cells Ca^2+^ signals and Cl^−^ currents are larger during tunneling as compared to SOCE alone (Fig. 1D-E). They are also induced significantly faster arguing that tunneling enhances the speed of the response by delivering Ca^2+^ more efficiently to its target, in this case ANO1 (Fig. 1F-G). In sweat cells and in vivo in the sweat gland blocking tunneling using CaTAr inhibits Cl^−^ secretion (Fig. 7H-I) and sweat production (Fig. 8A-D).

In the model proposed in Figure 8E, tunneling would be predicted to modulate both the amplitude and duration of the agonist-induced Ca^2+^ rise, a conclusion supported by the signal observed when SOCE in blocked (Fig. 8E, No SOCE). The modulation of the initial Ca^2+^ response by tunneling would depend on multiple factors: 1. The rate of IP_3_ production and degradation as tunneling depends on high IP_3_ levels in the cytosol. 2. The density of ERPMCS as sites for SOCE. 3. The expression levels of IP_3_R and KRAP that is the density of licensed IP_3_Rs. 4. The levels of STIM1, Orai1, PMCA, and SERCA. And 5. The distances between SOCE clusters, IP_3_Rs, and distal effectors. The dependency on these multiple factors allows cells the flexibility to modulate the extent of tunneling to fit their physiological needs. Despite the limited cell types tested to date, we have some confirmation of these predictions. For example, Ca^2+^ tunneling in frog oocytes induces a 30-fold larger Cl^−^ current compared to SOCE alone ^12^; whereas in salivary gland cells it is 1.5-fold higher, and in NCL cells it is at least 2.4-fold higher. In addition to modulating the amplitude and duration of Ca^2+^ signals, Ca^2+^ tunneling has also been shown to regulate the frequency of Ca^2+^ signals where it favors tonic over oscillatory Ca^2+^ transients ^15^.

Collectively our data show that store depletion remodels the Ca^2+^ signaling machinery in the cell cortex into subdomains both laterally in the plane of the ER and PM and axially within the cortical ER to support Ca^2+^ tunneling in delivering Ca^2+^ entering the cell through SOCE to distal effectors. This tunneling mechanism is important functionally in activating Cl^−^ secretion and sweat production.

## METHODS

### Cell culture and solutions

Hela cells were cultured in DMEM media containing 10 % fetal bovine serum (FBS) supplemented with penicillin (100 units.ml^−1^) and streptomycin (100 µg.ml^−1^). The cells were plated 24h before transfection on poly-lysine coated glass-bottom dishes (MatTek, U.S.A). NCL-SG3 cells were a gift from Stefan Feske (New York University) and were cultured in Williams E Media supplemented with 5 % FBS, with penicillin (100 units.ml^−1^), streptomycin (100 µg.ml^−1^), glutamine (4 mM), insulin (10 mg.l^−1^), transferrin (5.5 mg.l^−1^), selenium (6.7 µg.l^−1^), hydrocortisone (10 mg.l^−1^), epidermal growth factor (20 µg.l^−1^). For live cells experiments, cells were perfused using a peristaltic pump (Gilson Minipuls) at the speed of 1 ml.min^−1^. The standard saline contained (in mM) 145 NaCl, 5 KCl, 2 CaCl_2_, 1 MgCl_2_, 10 Glucose, 10 HEPES, pH 7.2, for Ca^2+^-free experiments, the Ca^2+^ was exchanged equimolarly with Mg^2+^.

### Plasmids and transfection

Transfection was performed using Lipofectamine 2000 (Thermo Fisher) according to the manufacturer’s instructions. mCherry-STIM1 and GFP-Orai1 were a gift from Rich Lewis (Stanford University, USA), EGFP-rIP3R1 from Colin Taylor (Cambridge University, UK), GFP–Mapper from Jen Liou (UT Southwestern, USA), and Ano1-GFP from Karl Kunzelmann (Regensburg, Germany). EGFP-hPMCA4b ^69^ and the Cl^−^ reporter mbYFPQS ^63^ were obtained from Addgene (#47589 and #80742) and phospholamban-GFP from Origene (#RG202712). CaTAr1 and 2 were custom made by Genewiz.

### Intracellular Ca^2+^ Imaging

To image cytoplasmic Ca^2+^ cells were loaded for 30 min at room temperature with either 2 µM Calbryte 590 AM (AAT Bioquest), Fluo4-AM or Fura 2-AM (Molecular Probes); 2 mM stocks were made in 20 % pluronic acid/DMSO. Imaging was performed on a Zeiss LSM880 confocal system fitted with a 40x/1.3 oil immersion objective using an open pinhole at a frame rate of 0.1 Hz. The following parameters were used: for Calbryte, λ_ex_=561 nm and λ_em_=566-679 nm; for Fluo4, λ_ex_=488 nm and λ_em_=493-574 nm. The expression level of either PLN-Map-GFP or SLN-Map-Ch were recorded using z-stacks and a pinhole set to 1AU cells at the beginning of the experiment using the following parameters: λ_ex_=488 nm and λ_em_=493-574 (GFP) and λ_ex_=561 nm and λ_em_=578-696 nm(mCherry). For Fura2 imaging of NCL cells was performed using a PTI Easy Ratio Pro system (software version 1.6.1.0.101; Horiba Scientific) composed of a DeltaRAMX monochromator and a CoolSnapHQ^2^ camera attached to an Olympus IX71 inverted microscope fitted with a 20x/0.75 lens. The cells were loaded for 30 min with 2 mM Fura2-AM in a Ca^2+^–containing media (composition below) at room temperature. Excitation was performed at 340 nm and 380 nm for a duration of 100 msec at a 0.1 Hz frame rate and the ratio of the fluorescence intensity at 340/380 measured.

### Airyscan Imaging

High resolution images were acquired using the Airyscan detector of a Zeiss LSM880 confocal microscope using the super-resolution mode (SR) and default image processing parameters. The 488 and 561 laser lines and a 488/561 MBS were used, and the emitted light recorded using either a single filter BP495-550/LP570 and a sequential line recording mode or a dual filter protocol (BP495-550/LP570 and BP420-480+495-550) and alternating z-stacks between both wavelengths. Z-stacks were recorded at the recommended intervals (typically 0.18 µm).

### TIRF Imaging

TIRF images were acquired on a AxioObserver Z1 microscope (Zeiss) using a 63x/1.46 lens at maximum angle and using the following parameters: for Alexa 488 and GFP: λex=488nm and λem=510/555; for Alexa 555 and mCherry λex=561 nm and λem=581/679.

### Acinar cell isolation and imaging

SMG acinar cells were enzymatically isolated from 2–to 4-month-old, C57BL/6J mice of both sexes. To isolate acinar cells, glands were extracted, connective tissue was removed, and glands were minced. Cells were placed in oxygenated dissociation media at 37°C for ∼30 min with shaking. Dissociation media consisted of Hank’s Balanced Salt Solution containing CaCl_2_ and MgCl_2_ (HBSS), bovine serum albumin (0.5%), and Collagenase Type II (0.2 mg/mL, Worthington). Cells were washed twice in HBSS with 0.5% BSA and resuspended in a HBSS solution containing 0.5% BSA and 0.02% trypsin inhibitor. Cells were then resuspended in imaging buffer (in mM) 10 HEPES, 1.26 CaCl_2_, 137 NaCl, 4.7 KCl, 5.5 glucose, 1 Na_2_HPO_4_, 0.56 MgCl_2_, at pH 7.4; with 5 μM FURA 2-AM and seeded onto a Cell-Tak coated coverslip to allow attachment of cells. Cells were then perfused with imaging buffer and stimulated with agonist. Ca^2+^ imaging was performed using an inverted epifluorescence Nikon microscope with a 40 X oil immersion objective (NA=1.3). Cells were alternately excited at 340 and 380 nm, and emission was monitored at 505 nm. Images were captured with a digital camera driven by TILL Photonics software. Image acquisition was performed using TILLVISION software.

### Patch clamp electrophysiology

For measurements of Cl^−^ currents, acinar cells were allowed to adhere to Cell-Tak coated glass coverslips for 15 min before experimentation. Coverslips were transferred to a chamber containing extracellular bath solution (in mM) 155 tetraethylammonium chloride to block K+ channels, 2 CaCl_2_, 1 MgCl_2_, 10 HEPES, pH 7.2. Ca2+ free bath solution substituted 1 mM EGTA for CaCl_2_. Cl^−^ currents in individual cells were measured in the whole-cell patch clamp configuration using pClamp 9 and an Axopatch 200B amplifier (Molecular Devices). Recordings were sampled at 2 kHz and filtered at 1 kHz. Pipette resistances were 3–5 MOhms, and seal resistances were greater than 1 GOhm. Pipette solutions (pH 7.2) contained (in mM) 60 tetraethylammonium chloride, 90 tetraethylammonium glutamate, 10 HEPES, 1 HEDTA (N-(2–hydroxyethyl) ethylenediamine-N,N’,N’ –triacetic acid) and 100 nM free Ca^2+^ were used to mimic physiological buffering and basal [Ca^2+^]_i_ conditions. Free [Ca^2+^] was estimated using Maxchelator freeware. Agonists were directly perfused onto individual cells using a multibarrel perfusion pipette.

### Sweat test

Sweating was measured using the starch/iodine technique ^64^. C57Bl6 male mice 3–to 5-month–old were used. The animals were injected with a control adenovirus (Ad-GFP, #1060, Vector Biolabs) or and adenovirus expressing CatAr1 (Ad-CMV, Vector Biolabs). The viruses in sterile PBS were injected: 50 µl at a titer of 10^8^ PFU/ml in the right hind foot pad of an anesthetized mice. The virus was allowed to express for 5 days prior to the sweat test. Images were acquired using an ERc5s Axiocam mounted on a STEMI 305 stereo microscope (Zeiss). Paw images were taken every 3 min and analyzed using ImageJ. The area covered by the black dots measured after 15 min was used to report sweating levels. Fluorescence images of the foot pads were taken using a fluorescence stereo microscope (Zeiss Lumar V12) equipped with a color camera (Zeiss Axiocam MR5).

### Immunocytochemistry

The cells were plated on glass bottom dishes and fixed using PFA (4%, 10 min), permeabilization was achieved using Triton x100 (10 min, 0.3%) and saturation for 1h using a mix of 10% goat serum and 1% bovine serum albumin. Primary antibody incubation was performed at 4°C overnight. The following primary antibodies were used at a 1:500 dilution: SERCA2 (NB300-581, Novus Biologicals), STIM1 (5668S, Cell Signaling and MAI-19451, Fisher), KRAP (14157-1-AP, Fisher), NFAT1 (5861S, Cell Signaling), ANO1 (14476, Cell Signaling). For IP_3_R1 detection a custom made monoclonal antibody targeting the following peptide: RIGLLGHPPHMNVNPQQPA (ProSci) was used ^70^. Secondary antibodies were anti-mouse and anti-rabbit coupled to either Ax488 or Ax555 (Molecular Probes) and used at a 1:2000 dilution at room temperature for 2 hours.

### Serial EM and 3D reconstruction

To prepare samples for TEM, Jurkat cell pellets were treated with a modified Karmovsky’s fix and followed by a secondary fixation in reduced osmium tetroxide as described previously ^71^. The samples were then En bloc stained with uranyl acetate and dehydrated with graded ethanol before being embedded in an Epon analog resin.

For serial EM sectioning the methods were as used previously ^72^. Briefly, 50 nm thick sections were collected on kapton tape using an ATUM (automated tape collecting microtome), then imaged by back scatter in a scanning electron microscope. Images were aligned in FIJI/Image J and further aligned in Reconstruct (version 1.1, https://synapseweb.clm.utexas.edu/software-0)^73^. The outlines of the cortical ER, and PM were hand-drawn by three different operators. Manual tracing and 3D renditions were performed in Reconstruct using Boissonnat Surface option for 3D representation.

### Data analysis and statistics

The imaging data was quantified using FIJI/ImageJ 1.51n ^74, 75^ and ZenBlue 2.3 (Zeiss). 3D reconstruction were performed using Imaris 9.5 (Bitplane). NND analysis was performed using the DiAna plugin ^76^ and colocalization using EzColocalization ^77^. The patch-clamp data was analyzed with Clampfit 10.0 (Molecular Devices). Statistics and data analysis were performed using Graphpad Prism 10.0.1 (GraphPad U.S.A). Values are given as Mean ± S.E.M and statistics were performed using either paired or unpaired Student’s t-test or ANOVA followed by Tukey’s test for multiple comparisons. P-values are ranked as follows * p≤0.05, ** p≤0.01, *** p≤0.001, **** p≤ 0.0001.

## Acknowledgments

We are grateful to the Imaging and Vivarium Cores at Weill Cornell Medicine Qatar (WCMQ) for their support. This work as well as the Cores are supported by the Biomedical Research Program at WCMQ (BMRP), a program funded by Qatar Foundation. We are grateful to colleagues who contributed clones and reagents as listed in the Methods section. We are also thankful to Jen Liou for helpful discussions during the early stages of this work regarding tunneling inhibition, and to Sandip Patel for suggesting the acronym CaTAr for the tunneling inhibitor during a presentation at a European Ca^2+^ Society (ECS) meeting.

## SUPPLEMENTARY INFORMATION

### Supplementary figure legends

**Supplemental Figure 1:**
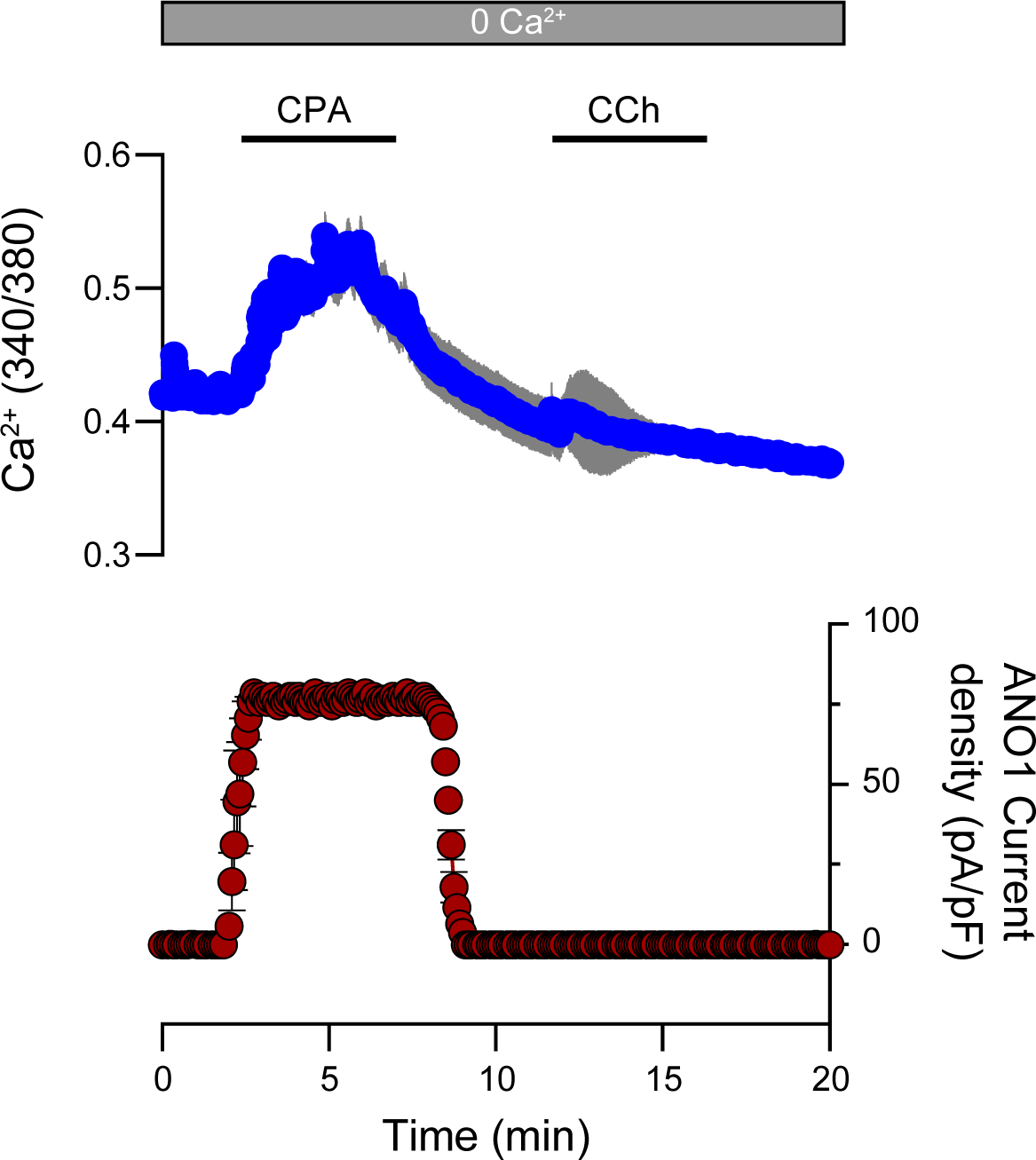
Store depletion in primary salivary gland cells. The transient application of CPA (30 µM) depletes ER Ca^2+^ stores as indicated by the rise in cytosolic Ca^2+^ (blue) which activates the Cl^−^ current (red). Following the washout of CPA, the application of carbachol (CCh, 10 µM) fails to elicit a response, indicating the effective depletion of the ER stores (n=4-8).

**Supplemental Figure 2:**
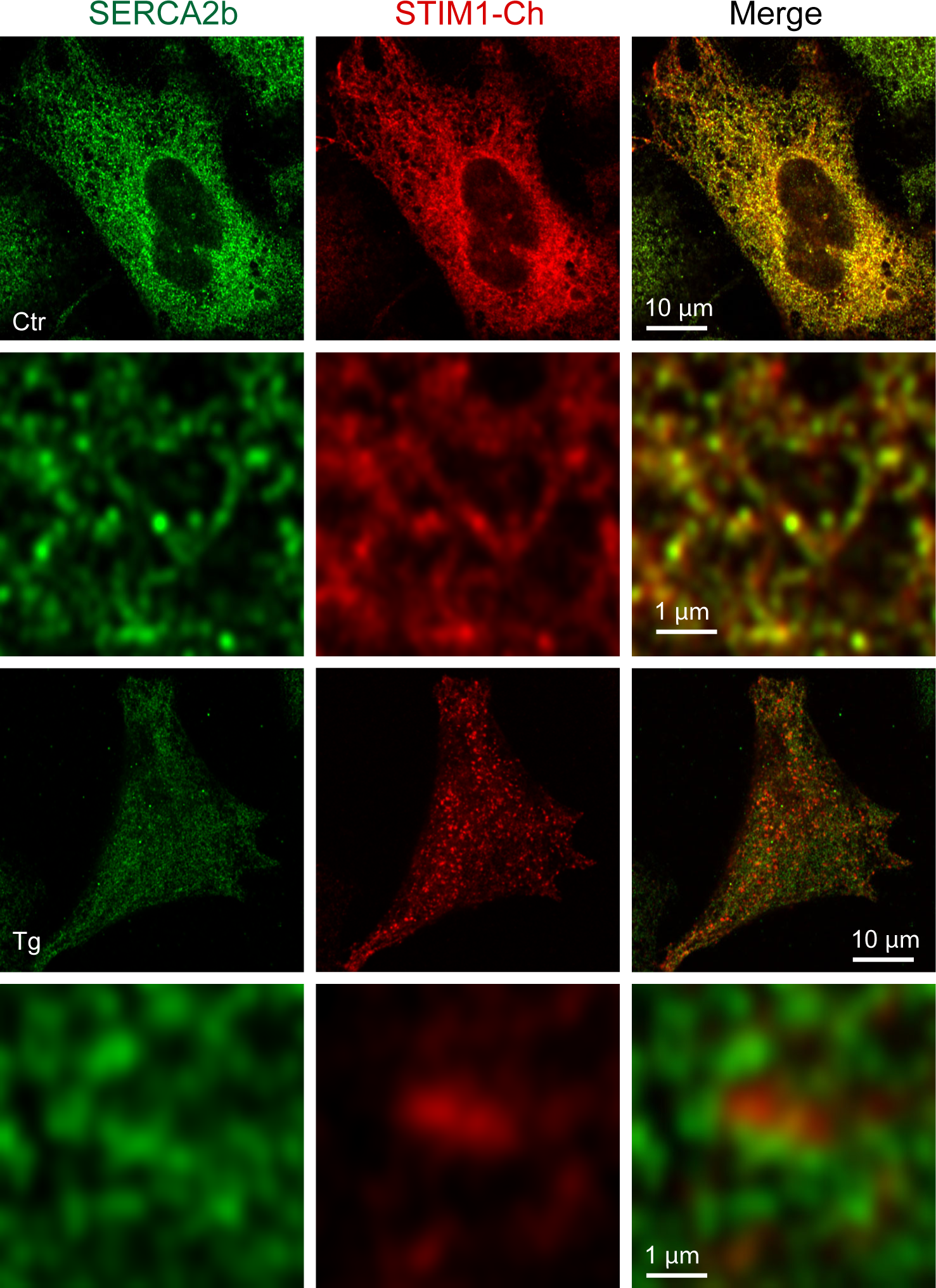
Localization of SERCA2b and mCh-STIM1. HeLa cells transfected with mCh-STIM1 and stained using a SERCA2b antibody. AiryScan images were taken before (Ctr) and after store depletion (Tg). Control images are acquired at a focal plane located in the middle of the cell and store-depleted images at the plasma membrane plane, where the STIM1 clusters localize. At rest STIM1 and SERCA2b colocalize in the ER cisternae, while after store depletion they separate isolate from each other at the PM focal place.

**Supplemental Figure 3:**
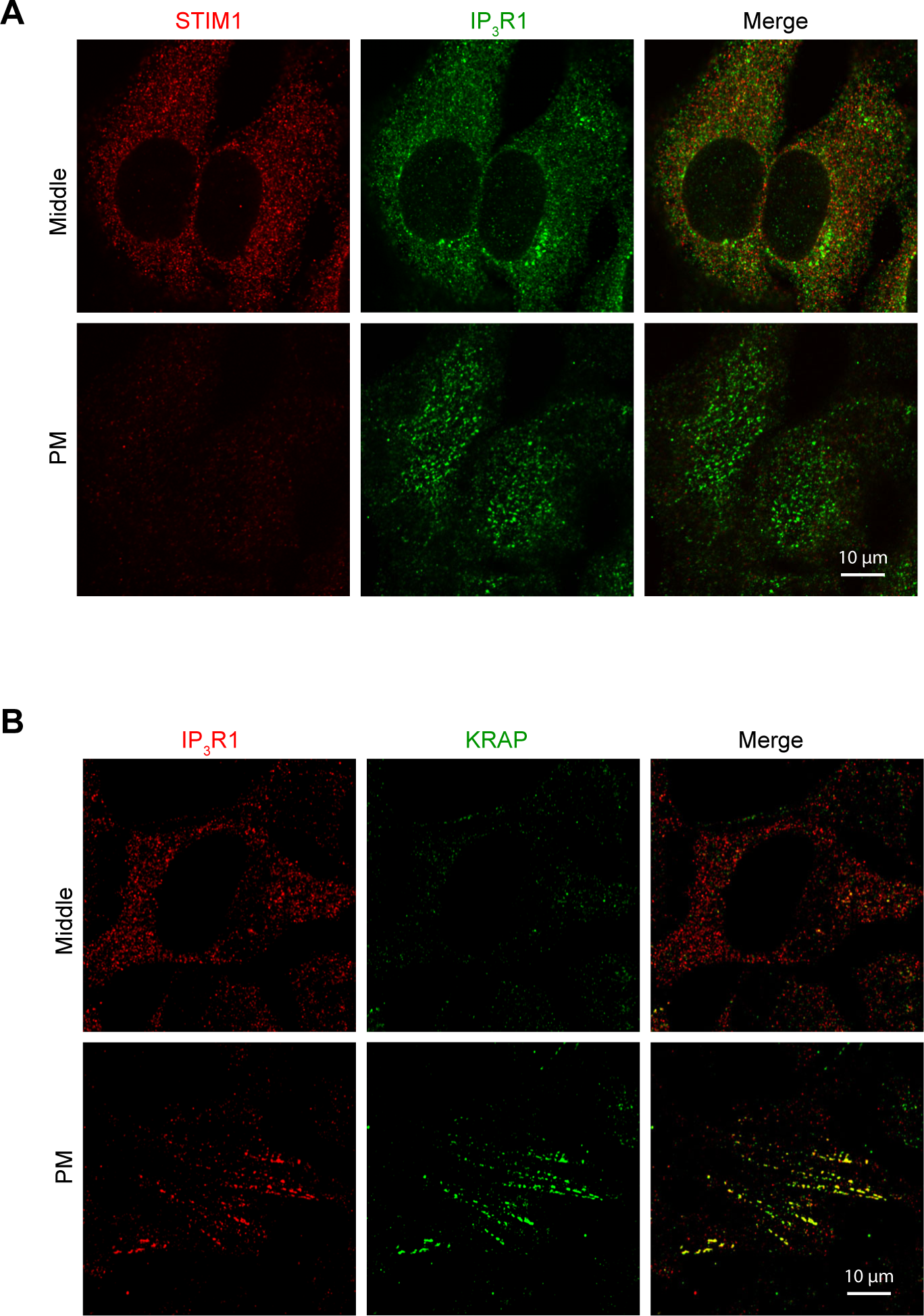
Localization of endogenous IP_3_R1, STIM1 and KRAP. (**A**) Relative localization of STIM1 and IP_3_R1 detected by immunofluorescence in HeLa cells at rest. While both proteins share the same intracellular compartment, there is no overlap of the signals at the PM where the IP_3_R1 fluorescence reveals the “licensed” receptors. **(C)** Co–localization at the PM focal plane of IP_3_R1 and KRAP in HeLa cells at rest.

**Supplemental Figure 4:**
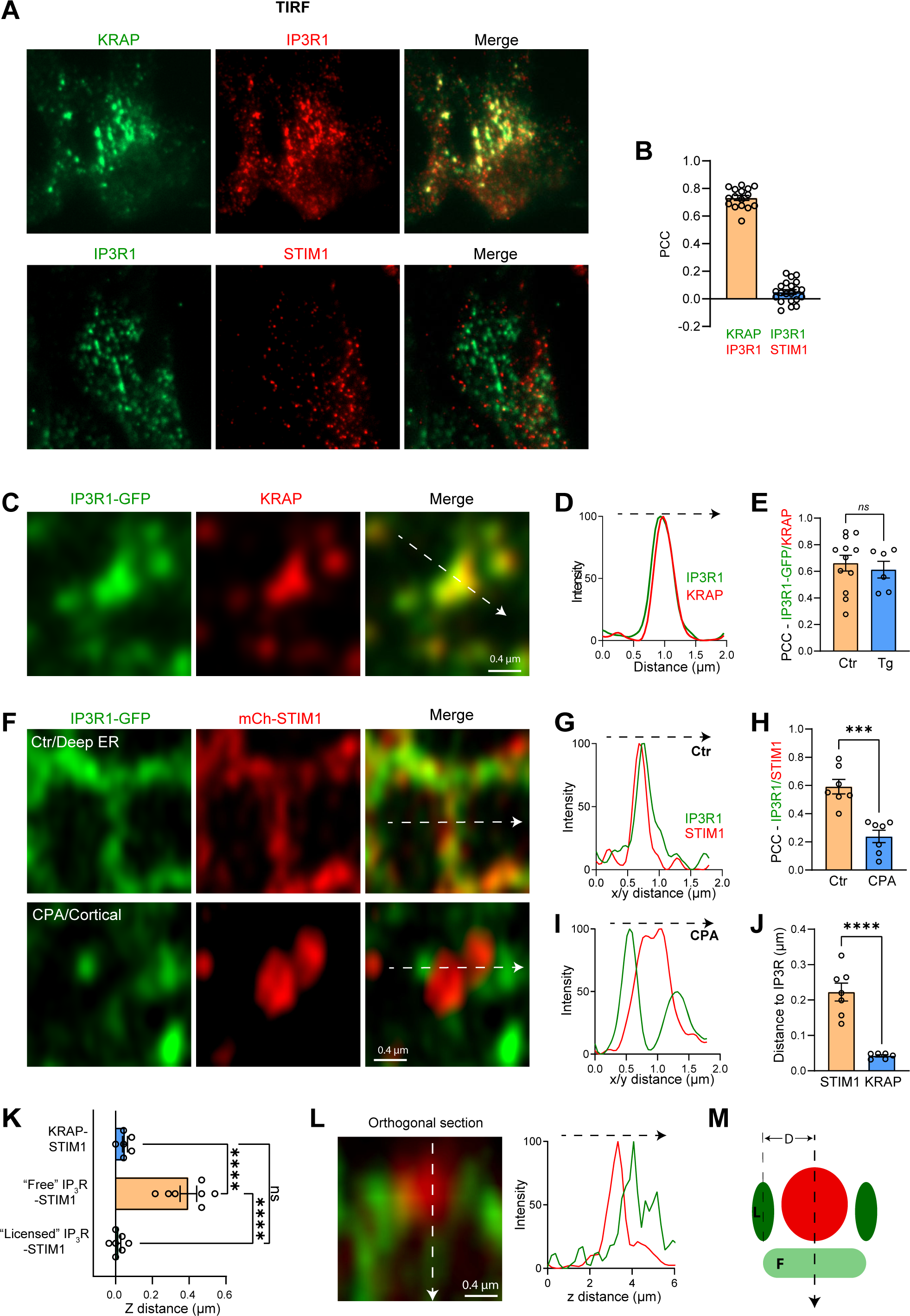
Localization of IP_3_R1, STIM1, and KRAP. (**A**) TIRF images of STIM1, KRAP, and IP_3_R1 detected by immunofluorescence in HeLa cells after store depletion. **(B)** Bar chart summarizing colocalization (PCC) of IP_3_R1 with KRAP but not with STIM1 (n=16-21; unpaired t-test). **(C)** Airyscan images or “licensed” IP_3_R1-GFP and KRAP detected by immunofluorescence at the PM plane (n=6-12; unpaired t-test). **(D)** Relative intensities measured along the line indicated by the white arrow in (C). **(E)** Colocalization (PCC) between KRAP and IP_3_R1-GFP before (Ctr) and after store depletion (Tg). **(F)** Airyscan images of IP_3_R1-GFP and mCh-STIM1 inside the cell (deep ER, in control conditions) and at the PM after store depletion (CPA/Cortical). **(G)** Relative intensities measured along the line indicated by the white arrow in (F) in cells at rest. **(H)** Colocalization (PCC) of IP_3_R1-GFP and mCh-STIM1 before and after store depletion (CPA) (n=7; paired t-test). **(I)** Relative intensities along the line indicated by the white arrow in (F) in store depleted cells. **(J)** Lateral distance between the mCh-STIM clusters and either IP_3_R1-GFP or endogenous KRAP after store depletion (n=6-7; unpaired t-test). **(K)** Axial distance between STIM1 clusters after store depletion and KRAP, “Free” IP_3_R1-GFP, or “Licensed” IP_3_R1-GFP, as indicated. “Free” IP_3_R1 are receptors that localized deeper in the cell and do not colocalize with KRAP. **(L)** Example of an orthogonal section through a mCh-STIM1 cluster highlighting the localization of “licensed” IP_3_R1-GFP outside the cluster and “Free” IP_3_R deeper within the ER away from the STIM1 cluster as illustrated by the intensity plot (white arrow). **(M)** Cartoon summary of the distribution of STIM1 (red) after store depletion relative to “Licensed” (indicated as L) and “Free” (indicated as F) IP_3_R1-GFP. The lateral distance between STIM1 clusters and licensed IP_3_R1 as measured in panel J is indicated as D.

**Supplemental Figure 5:**
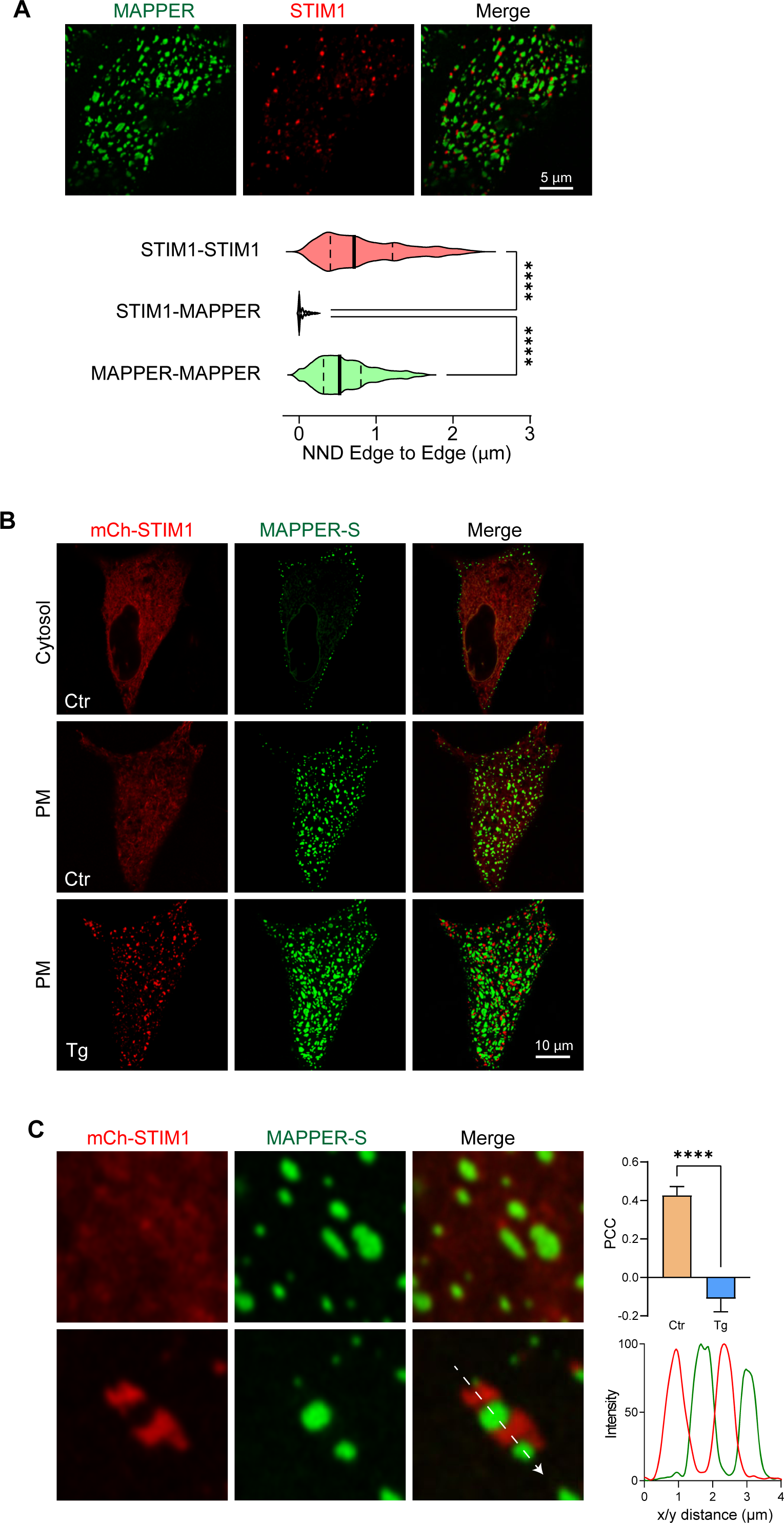
Localization of MAPPER-S relative to STIM1. (**A**) Airyscan images of MAPPER-GFP (MAPPER) and mCh-STIM1 following store depletion with CPA in HeLa cells showing that they do not colocalize. The violin plots indicate the Nearest Neighbor Distance (NND) between the edges of the STIM1 and of the MAPPER clusters (n=626–1585, outliers removed using the ROUT routine, one-way ANOVA). **(B)** Relative localization of mCh-STIM1 and short MAPPER (MAPPER-S) in control conditions (Ctr) and after store depletion (Tg) at the whole cell level. **(C)** High magnification images of mCh-STIM1 and MAPPER-S clusters with the corresponding PCC (n=11, paired t-test) and line scan profile measured along the white arrow.

**Supplemental Figure 6:**
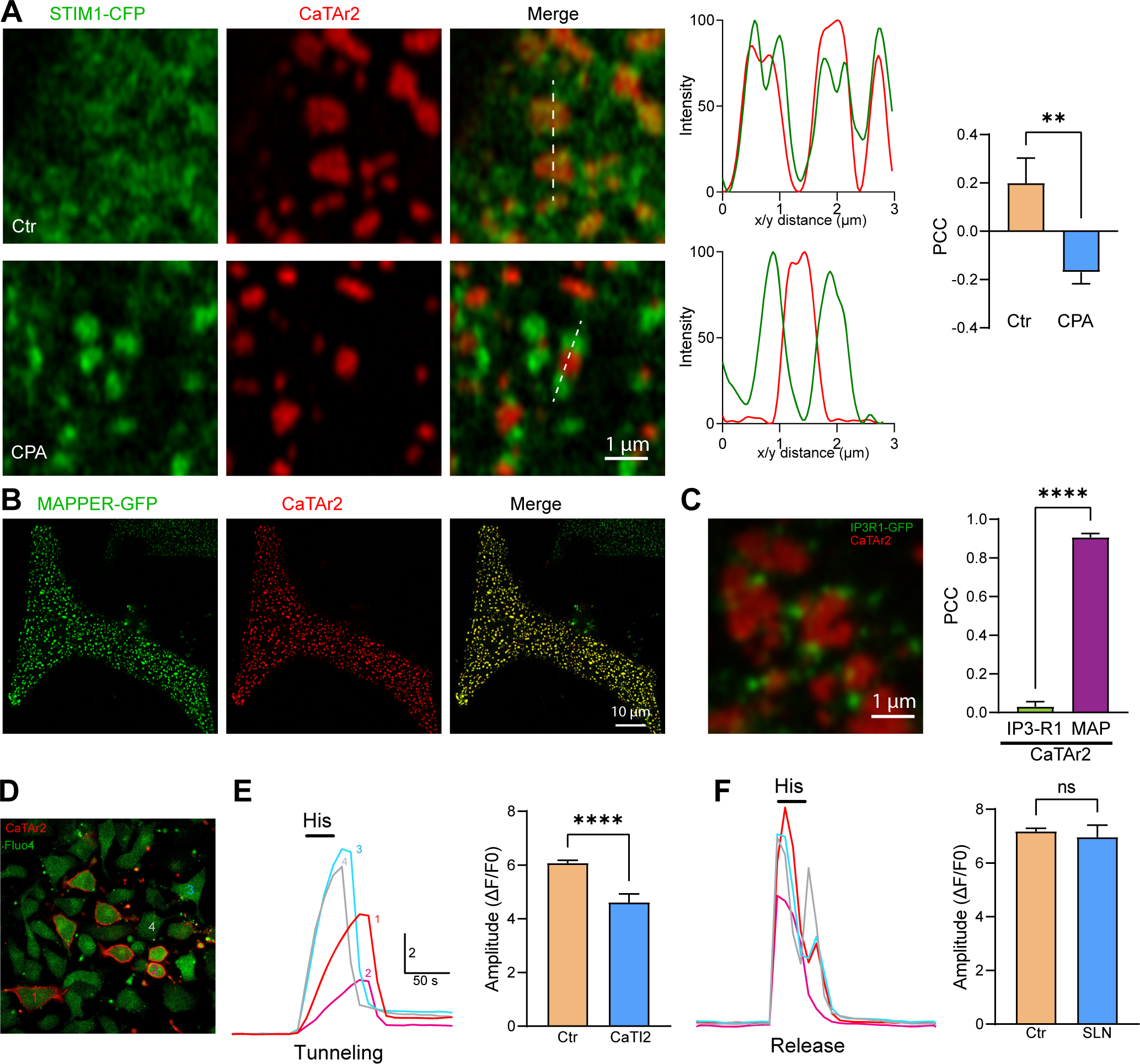
CaTAr2. (**A**) Localization of CaTAr2 relative to STIM1-CFP before (Ctr) and after store depletion (CPA). Colocalization analysis using either intensity plots along the white line or PCC measurements confirm that CaTAr2 localizes in a similar fashion to CaTAr1 and MAPPER relative to STIM1 clusters (n=5-10; unpaired t-test). **(B)** Colocalization of MAPPER-GFP and CaTAr2 at the whole cell level. **(C)** Localization and PCC between CaTAr2 and IP_3_R1-GFP. The colocalization of CatAr2 with MAPPER was used as a reference value for the PCC analysis (n=6-14; unpaired t–test). **(D)** Confocal images of HeLa cells expressing CaTAr2 and loaded with the Ca^2+^ Fluo4-AM. The numbered cells refer to the traces in E and F. **(E)** Ca^2+^ tunneling traces obtained after store depletion with CPA indicate that cells expressing CaTAr2 have a lower tunneling capacity. The bar chart on the right summarizes the inhibition of Ca^2+^ tunneling by CaTAr2. **(F)** Ca^2+^ release traces obtained on the same cells as in E by applying histamine (His, 100 μM) after store refilling. The bar chart on the right summarizes the absence of effect of CaTAr2 on Ca^2+^ release (n=42–342; unpaired t-test).

**Supplemental Figure 7:**
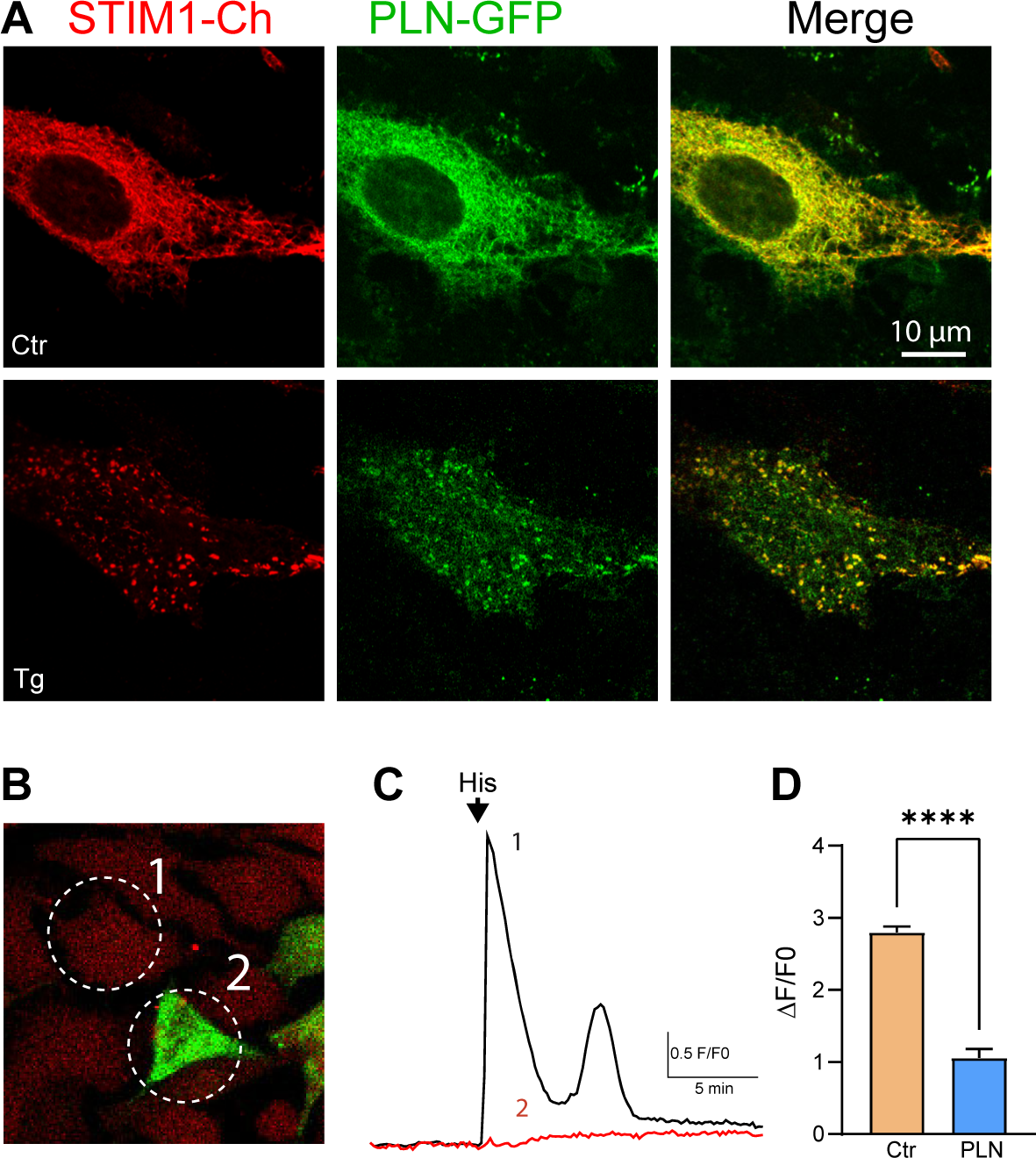
Phospholamban inhibits SERCA. (**A**) Localization of phospholamban-GFP (PLN-GFP) and mCh-STIM1 in HeLa cells before (Ctr) and after store depletion (Tg). **(B)** PLN-GFP transfection followed by loading cells with Calbryte 590. The numbered cells match the traces in C. **(C)** Ca^2+^ release induced by histamine (His, 100 μM) leads to Ca^2+^ release in cell#1 which does not express PLN-GFP but not in cell#2 which expressed high levels of PLN-GFP. **(D)** Bar chart summarizing the inhibition of Ca^2+^ release in PLN-expressing cells. The recordings were performed in the presence of 1 mM lanthanum ions in the extracellular media to inhibit a potential contribution of Ca^2+^ influx or recycling through the PM (n=38-176; unpaired t-test).

**Supplemental Figure 8:**
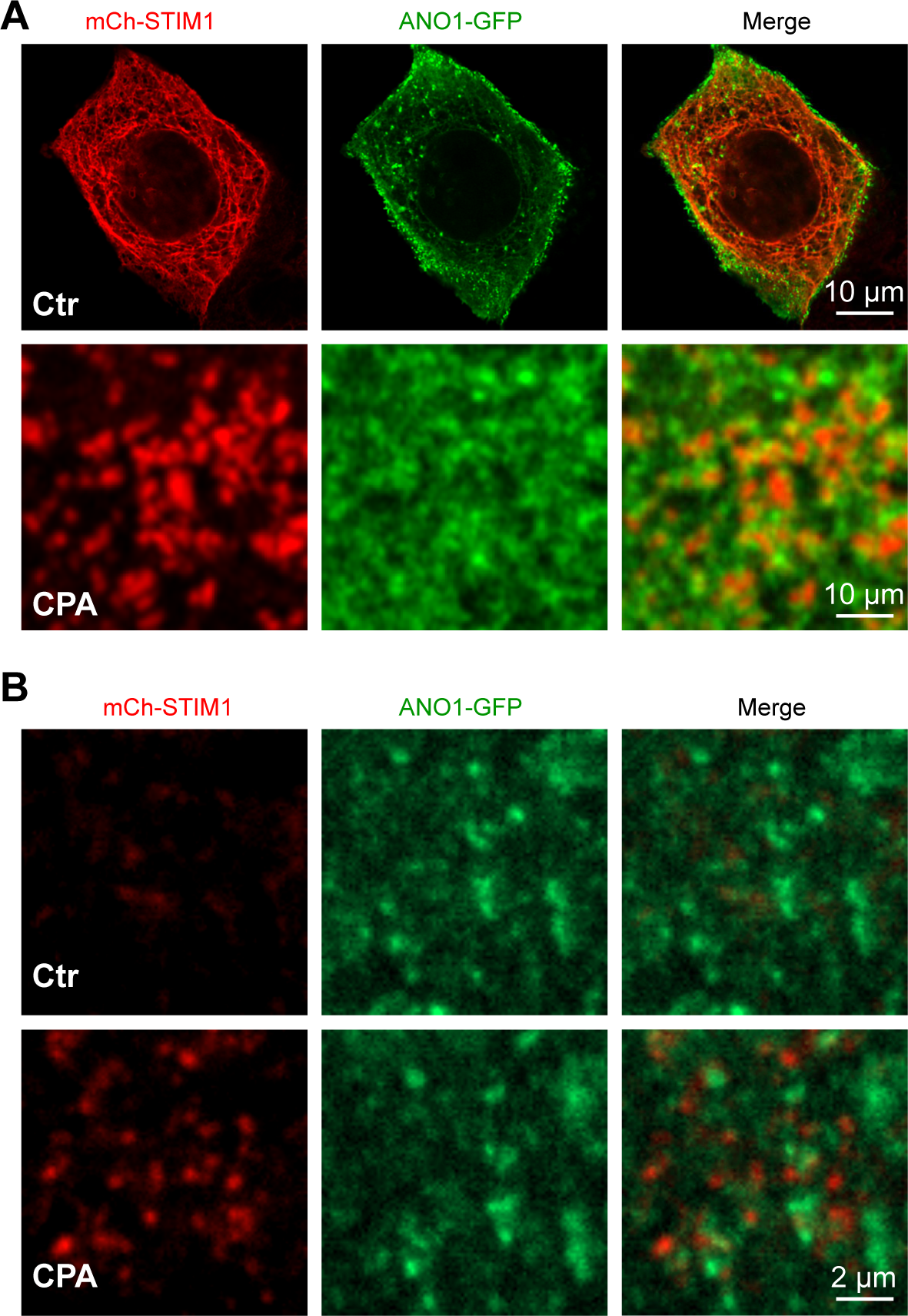
Localization of ANO1-GFP and mCh-STIM1. (**A**) The relative localization of ANO1-GFP and mCh-STIM1 in NCL cells after store depletion is similar to the endogenous ANO1 channel and suggests an separation of ANO1 from the STIM1 cluster. **(B)** TIRF imaging of ANO1-GFP and mCh-STIM1 during store depletion with CPA confirmed the separation of the two proteins at the PM. Ctr and CPA images are from the same region of interest.

**Supplemental Video 1: Formation of STIM1 clusters adjacent to MAPPER** A HEK293 cell overexpressing MAPPER-GFP (green) and mCh-STIM1 (red) is imaged over time using TIRF microscopy. Store depletion is induced using thapsigargin leading to the formation of STIM1 clusters next to MAPPER-GFP contact sites. The frame rate is 1.2 Hz for a total duration of 10 minutes.

**Supplemental Video 2: Formation of STIM1 clusters adjacent to CaTAr1** A HeLa cell overexpressing CaTAr1 (green) and mCh-STIM1 (red) is imaged over time using TIRF microscopy. Store depletion is induced using thapsigargin leading to the formation of STIM1 clusters next to CaTAr1 expression sites. The frame rate is 0.1 Hz for a total duration of 36 minutes.

